# Differences in the regulatory strategies of marine oligotrophs and copiotrophs reflect differences in motility

**DOI:** 10.1101/2022.07.21.501054

**Authors:** Stephen E. Noell, Elizabeth Brennan, Quinn Washburn, Edward W. Davis, Ferdi L. Hellweger, Stephen J. Giovannoni

**Affiliations:** Department of Microbiology, Oregon State University, Corvallis, OR, USA; Center for Quantitative Life Sciences, Oregon State University, Corvallis, OR, USA; Water Quality Engineering, Berlin TU, Berlin, Germany

**Keywords:** regulation, copiotroph, oligotroph, chemotaxis, bacterioplankton, alanine, modeling

## Abstract

Aquatic bacteria frequently are divided into lifestyle categories *oligotroph* or *copiotroph*. Oligotrophs have proportionately fewer transcriptional regulatory genes than copiotrophs and are generally non-motile/chemotactic. We hypothesized that the absence of chemotaxis/motility in oligotrophs prevents them from occupying nutrient patches long enough to benefit from transcriptional regulation. We first confirmed that marine oligotrophs are generally reduced in genes for transcriptional regulation and motility/chemotaxis. Next, using a non-motile oligotroph (*Ca*. Pelagibacter st. HTCC7211), a motile copiotroph (*Alteromonas macleodii* st. HOT1A3), and [^14^C]L-alanine, we confirmed that L-alanine catabolism is not transcriptionally regulated in HTCC7211 but is in HOT1A3. We then found that HOT1A3 took 2.5-4 min to initiate L-alanine oxidation at patch L-alanine concentrations, compared to <30s for HTCC7211. By modeling cell trajectories, we predicted that, in most scenarios, non-motile cells spend <2 min in patches, compared to >4 mins for chemotactic/motile cells. Thus, the time necessary for transcriptional regulation to initiate prevents transcriptional regulation from being beneficial for non-motile oligotrophs. This is supported by a mechanistic model we developed, which predicted that HTCC7211 cells with transcriptional regulation of L-alanine metabolism would produce 12% of their standing ATP stock upon encountering an L-alanine patch, compared to 880% in HTCC7211 cells without transcriptional regulation.

## Introduction

Heterotrophic marine bacteria play a major role in the fate of oceanic organic carbon (Hansell *et al*., 2009) by respiring as much as 50% of ocean primary productivity (Azam *et al*., 1983). These organisms employ different life strategies to exploit varied opportunities for nutrient acquisition, influencing their success in harvesting dissolved organic matter (DOM) resources. One common way of categorizing these life strategies is *copiotrophic* or *oligotrophic*, with microorganisms existing on a spectrum between these extremes. Copiotrophs have been characterized as “feast or famine” bacteria - they are capable of large changes in growth rate in response to substrate availability (Koch, 1971). Oligotrophs are adapted to live off low nutrient concentrations and thus make only small adjustments in growth rates in response to changes in nutrient concentrations (Poindexter, 1981). These lifestyle strategies are generally associated with physiological adaptations, especially in marine microbes: copiotrophs are generally larger in cell size than oligotrophs, have larger genomes with higher GC content, and are generally motile (Lauro *et al*., 2009; DeLong *et al*., 2010; Kempes *et al*., 2012, 2016; Giovannoni *et al*., 2014; Chuckran *et al*., 2021; Weissman *et al*., 2021), although these associations are debated (Westoby *et al*., 2021).

A fundamental difference between oligotrophs and copiotrophs that remains relatively unexplored is the reduced transcriptional regulation of carbon uptake and oxidation functions in oligotrophs, which has been shown both on the genomic and physiological level (Lauro *et al*., 2009; Cottrell and Kirchman, 2016; Sun *et al*., 2016; Weissman *et al*., 2021). This is not to say that oligotrophic marine bacteria completely lack transcriptional regulatory mechanisms. In the abundant marine heterotrophic bacterial group SAR11, several studies have documented low amplitude diel transcriptional changes in a variety of genes and strong transcriptional responses to iron, sulfur, phosphorous, and nitrogen starvation, which are common causes of environmental stress that have corresponding transcriptional regulators in these cells (Smith *et al*., 2010, 2013, 2016; Ottesen *et al*., 2014; Carini *et al*., 2015). Oligotrophs seem to have evolved reduced transcriptional regulation of carbon uptake and oxidation functions, while retaining post-transcriptional regulatory mechanisms, such as riboswitches, and kinetic regulatory mechanisms (Giovannoni, 2017). In principle, these post-transcriptional mechanisms respond more rapidly to change and require less coding capacity to implement than transcriptional regulation but are not as versatile.

The ocean environment on the microscale is heterogeneous, both in coastal regions and the oligotrophic open ocean (Pilskaln *et al*., 2005; Kostadinov *et al*., 2009; Stocker, 2012), with extreme gradations of nutrient levels in micro-patches, such that the exposure of marine bacterioplankton to nutrients is highly variable (Stocker, 2012). Oligotrophs and copiotrophs, due to differences in motility and chemotaxis (Lauro *et al*., 2009), experience ocean microscales in vastly different ways. Copiotrophs are adapted to maximize their time spent in these nutrient patches, primarily by attaching to particulate organic matter or using chemotaxis and motility to identify, move towards, and maximize time spent in ephemeral nutrient patches (Poindexter, 1981; Lambert *et al*., 2019). Oligotrophs inhabit the same patchy environments as copiotrophs (Kostadinov *et al*., 2009; Stocker, 2012) but have fewer encounters with patches and particles than chemotactic cells due to their general lack of motility and chemotaxis (Blackburn *et al*., 1998), which is partially attributable to their small cell sizes (Mitchell, 1991; Dusenbery, 1997).

To explain differences in the regulation of carbon metabolism between oligotrophs and copiotrophs, we hypothesized that the lack of chemotaxis and motility in oligotrophs limits their time in patches of nutrients to periods too short for transcriptional regulatory systems to initiate and be effective (generally ~3 minutes (Pardee and Prestidge, 1961; Kepes, 1963)). The chemotactic ability of copiotrophs in principle allows cells to stay in patches long enough for transcriptional regulatory systems to be fully induced so that cells can reap the benefits of enhanced gene expression. We reasoned that oligotrophs, at the whim of the environment, might rely more on faster-acting and lower amplitude post-transcriptional regulation to take advantage of whatever patches they encounter.

We explored this hypothesis in several steps. We began by using a recently proposed quantitative definition of oligotrophy/copiotrophy based on predicted maximal growth rates to provide a clearer resolution to the two assumptions of our hypothesis: do marine oligotrophs have reduced genomic proportions of transcriptional regulators and chemotaxis/motility genes? Next, we developed an experimental system to provide quantitative measurements to complement later modeling efforts. Our experimental system was comprised of a copiotroph, *Alteromonas macleodii* st. HOT1A3 (HOT1A3 throughout; an open-ocean, motile gammaproteobacteria with complex regulatory systems (Ivars-Martinez *et al*., 2008; Fadeev *et al*., 2016)), and an oligotroph, *Candidatus* Pelagibacter st. HTCC7211 (HTCC7211 throughout; open-ocean representative of the SAR11 clade – all known members are non-motile – with minimal regulation (Giovannoni, 2005, 2017)), and a metabolite (L-alanine) whose common metabolic pathway (conversion into pyruvate via L-alanine dehydrogenase, *ald* gene) and regulation have been well-characterized (Berberich *et al*., 1968; Jeong *et al*., 2015). We confirmed the presence/absence of the trait of interest (transcriptional regulation of metabolism) in each strain using radiotracer experiments, reverse transcription qPCR (RT-qPCR), and mechanistic modeling. Third, we measured the time between exposure to the nutrient and initiation of oxidation of that metabolite by each strain at a variety of nutrient concentrations. Finally, we compared these measured oxidation initiation times to simulated residence times of motile and non-motile cells in nutrient patches, concluding that transcriptional regulation of carbon assimilation pathways would not benefit many oligotrophic marine bacterioplankton because of their lack of motility/chemotaxis, and that “always on” catabolic systems represent a different lifestyle strategy that is advantageous for cells that are unable to direct their movement through the ocean microenvironment.

## Results and Discussion

### Exploration of Chemotaxis/Motility Genes in Marine Microbes

We began by providing confirmatory evidence for the two assumptions of our hypothesis: marine oligotrophs are reduced in transcriptional regulation and tend to be non-chemotactic/motile compared to copiotrophs. Several lines of prior evidence support our first assumption: 1) oligotrophs have fewer genes for transcriptional regulation, relative to genome size (Giovannoni, 2005; Lauro *et al*., 2009; Giovannoni *et al*., 2014; Held *et al*., 2019; Lambrecht *et al*., 2020; Graham and Tully, 2021; Weissman *et al*., 2021; Chiriac *et al*., 2022); 2) the proportion of protein coding genes transcribed and translated in oligotrophic cells is much higher than copiotrophs (Smith *et al*., 2010, 2013, 2016; Cottrell and Kirchman, 2016); 3) when individual carbon assimilation functions have been investigated experimentally in oligotrophs, in most cases they were found to be constitutively expressed (Sun *et al*., 2016; Giovannoni *et al*., 2019). In support of our second assumption, aquatic pelagic oligotrophs have been found previously to be reduced in motility genes (Lauro *et al*., 2009; Chiriac *et al*., 2022), although this result did not hold for RefSeq genomes across all habitats (Weissman *et al*., 2021).

We used a recently proposed quantitative definition of oligotrophy/copiotrophy based on predicted maximal growth rate (<5h were defined as copiotrophs, ≥5h were defined as oligotrophs) (Weissman *et al*., 2021). The use of this definition is not without issues, especially since growth rate is only one aspect of lifestyle strategy and oligotrophs/copiotrophs are known to differ in a variety of traits (e.g., (Chuckran *et al*., 2021)). However, its value for this study lies in its quantitative nature; cells can often be misclassified as oligotrophs or copiotrophs based solely on the environment they were isolated from or based on their growth characteristics in the lab under select nutrient conditions. We used metagenome assembled genomes (MAGs) from (Tully *et al*., 2018) with the same oligotroph/copiotroph categorizations generated in (Weissman *et al*., 2021). We found that these marine oligotrophic genomes have characteristics common to oligotrophs (small genome size and low %GC) (Figure 1A-B). As found previously, the oligotrophs were significantly (Mann–Whitney test with Benjamini–Hochberg correction, p-value ≤ 0.05) reduced in genes involved in transcriptional regulation compared to copiotrophs (Figure 1D). They were also significantly reduced in chemotaxis genes (Figure 1D), in contrast to the results of (Weissman *et al*., 2021). However, no significant difference was observed in motility genes (Figure 1D). A cell will benefit from non-directed motility by moving out of spaces where nutrients have been exhausted (Bondoc *et al*., 2016), but the significance of this mechanism has not been quantified.

**Figure 1.**
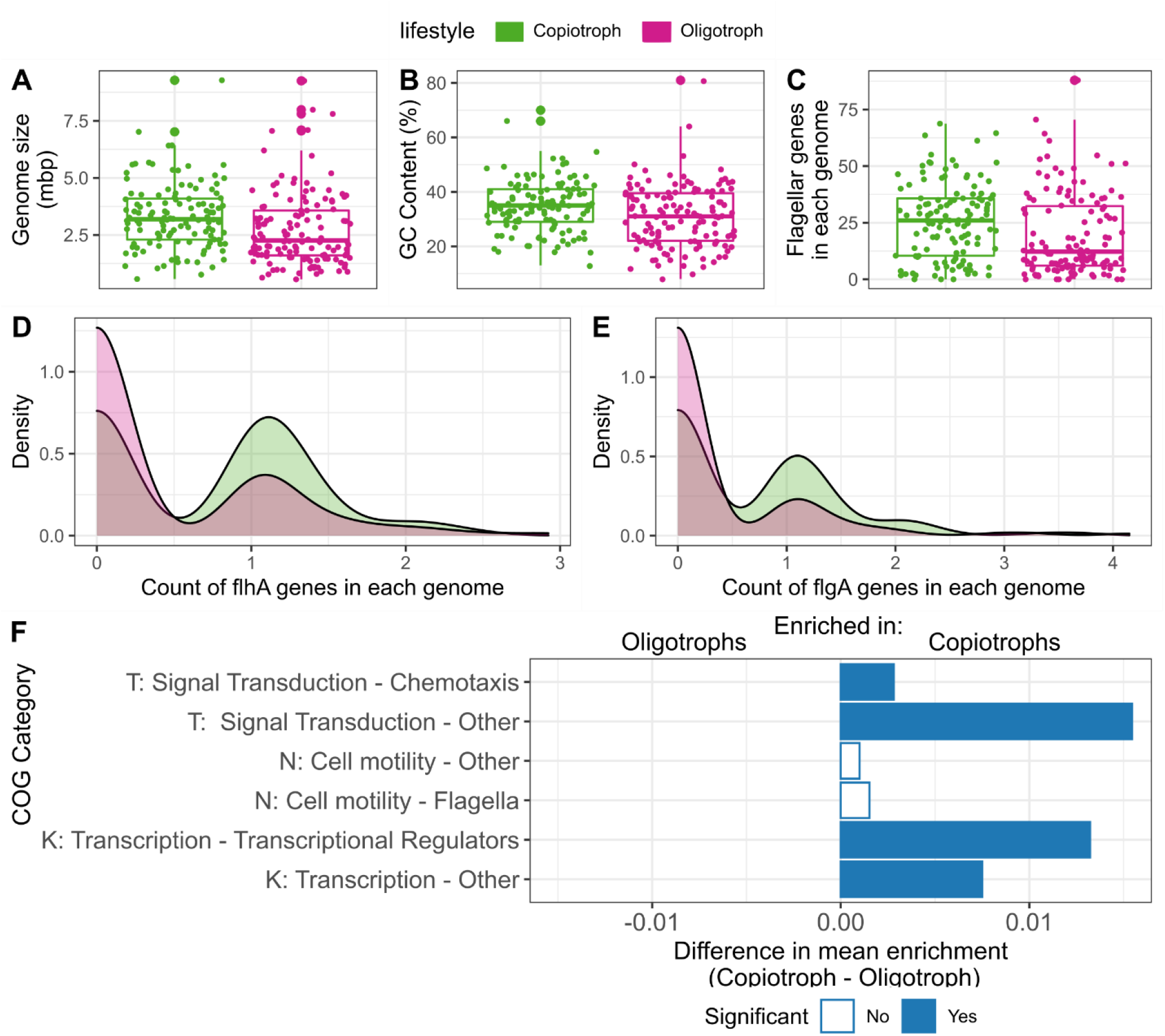
Comparison of genomic and functional characteristics of marine copiotroph/oligotroph metagenome assembled genomes (MAGs) from Tully et al. (Tully *et al*., 2018), with lifestyle strategy defined by predicted maximal growth rate based on codon usage patterns from Weissman et al. (Weissman *et al*., 2021). (A, B) Comparison of genome characteristics of the two lifestyle categories from this data set. (C) Comparison of the number of flagellar genes between copiotrophs and oligotrophs; each point represents the count of flagellar genes within one genome. Flagellar gene count was normalized by genome completeness, not genome size, to account for potentially missed flagellar genes. (D, E) Density distribution of the count of either (D) *flhA* or (E) *flgA* genes in copiotrophic or oligotrophic genomes; these genes are required for proper flagellar functioning. Copiotrophs are more likely to have one or more copies of these genes, while oligotrophs are more likely to have zero copies of these genes. (F) Enrichment of different, selected COG categories in copiotrophs or oligotrophs; enrichment is a proportion of the genes in a given COG category divided by the total number of genes across all COG categories. Category T was split between COG categories having “Chemotaxis” in their description and all others. Category N was split between categories with “Flagella” in their description and all others. Category K was split between categories with “Transcriptional Regulator” in their description and all others. Significant enrichment was tested using Mann– Whitney tests with a Benjamini–Hochberg correction (α=0.05); significant enrichment indicates an adjusted p-value ≤ 0.05.

We explored whether the smaller average genome size of oligotrophs was biasing the outcome for motility, given that flagellar functioning generally requires a certain threshold of genes for proper functioning (generally at least 20 genes, although some species have multiple flagella and thus over 100 flagellar genes (Liu and Ochman, 2007)). We found that copiotrophs had significantly larger numbers of flagellar genes per genome, with a median value above the ~20 gene threshold (median: 12.2 and 26 for oligotrophs and copiotrophs, respectively; p = 0.001, Mann– Whitney test, Benjamini–Hochberg correction) (Figure 1C). Two flagellar genes, *flhA* and *flgA*, are known to be required for proper flagellar operation (Rossmann and Beeby, 2018; Inoue *et al*., 2021); these genes were also significantly more likely to be found in copiotrophs than oligotrophs (p = 0.0001 and 7E-5 for *flhA* and *flgA*, respectively, Mann– Whitney test, Benjamini–Hochberg correction) (Figure 1D, E). These results, combined with the recent finding that chemotactic and flagellar genes have co-evolved in some bacterial groups (Mo *et al*., 2022), indicate that marine oligotrophs are generally non-chemotactic and non-motile.

### Experimental Observations of L-alanine Uptake and Metabolism: Confirmation of Traits

Next, we developed an experimental system to provide detailed experimental data to explore our hypothesized connection between regulatory strategy and motility/chemotaxis. We began by determining the presence/absence of our trait of interest (transcriptional regulation of LALA uptake and metabolism) in our two strains using physiological measures (uptake and oxidation of [^14^C]L-alanine ([^14^C]LALA)) and transcript levels (using reverse transcription quantitative real-time PCR (RT-qPCR)). For [^14^C]LALA uptake and metabolism measurements, we used a high initial concentration of LALA (4 μM) to simulate patch conditions (Pocklington, 1971; Lee and Bada, 1977; Mopper and Lindroth, 1982; Lu *et al*., 2014). Collected uptake and oxidation data is presented in Figure 2 (scatter plots; measured values are depicted in time course plots; experimental data is shown in dots, while the lines are model results that will be discussed subsequently), with calculated rates between time points depicted in the bar plots. A consistent rate of uptake or oxidation could indicate an absence of transcriptional regulation of the necessary genes, while an increase in uptake or oxidation rates could indicate the opposite.

**Figure 2.**
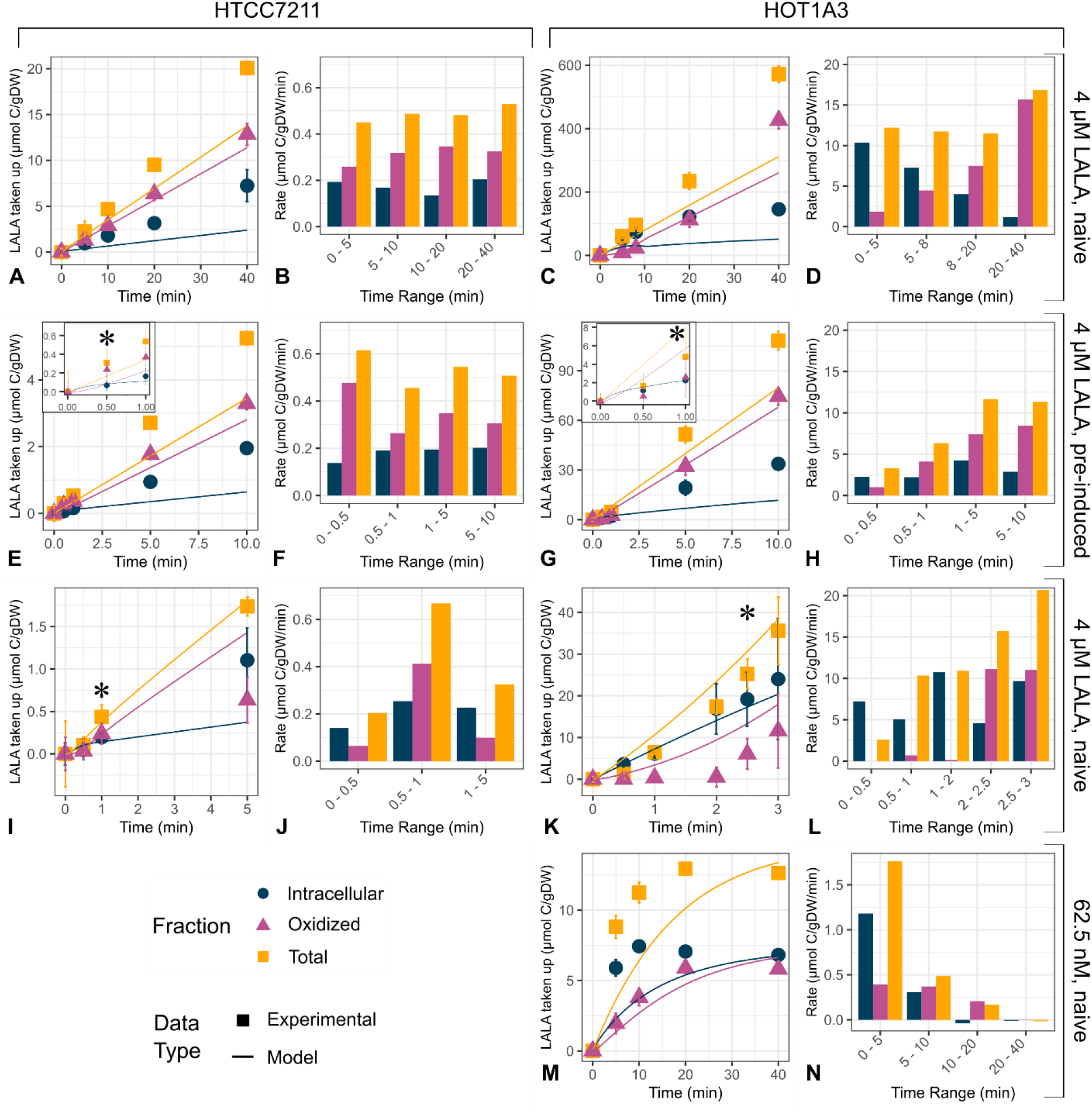
Uptake and oxidation of [^14^C]L-alanine ([^14^C]LALA) in (A-B, E-F, I-J) an oligotroph, *Ca. P*. st. HTCC7211, and (C-D, G-H, K-N) a copiotroph, *A. macleodii* st. HOT1A3, to determine induction and time to initiation of uptake and oxidation. (A, C, E, G, I, K, M) Time course of uptake and oxidation of [^14^C]LALA in both strains. Points are experimental data, while the lines are the output of the mechanistic model. (B, D, F, H, J, L, N) Bar plots of calculated [^14^C]LALA uptake, oxidation, and intracellular accumulation rates in the indicated time intervals based on data presented in time course plots. Cells were grown either without (A-D, I-N) or with (E-H) 4 μM LALA prior to harvesting, washing, and starving. Cells were then incubated with either 4 μM (A-L) or 62.5 nM (M-N) of [^14^C]LALA. (A-D) To determine presence of induction, time points out to 40 min were taken. (E-H) The experiment was repeated to 10 min with cells that were pre-exposed to LALA for 2 generations before cells were harvested. (I-L) To determine the time until oxidation initiation, short time points were used. For detailed explanation of the different fractions, see Methods. Error bars on points are the standard deviation of triplicate cultures. Where there are error bars, they are too small to be visible. * indicates oxidation initiation, the first time point where oxidation was greater than zero and the standard deviation did not overlap zero. Note that y- and x-axis scales differ between plots.

In initial uptake/oxidation experiments, the cells used were naïve (i.e., not exposed to LALA prior to the experiment). Over the initial 40-min experiment, in HTCC7211, LALA uptake and oxidation rates remained constant: 0.5 and 0.3 μmol C/g dry weight (DW)/min for uptake and oxidation rates, respectively (Figure 2A-B). In HOT1A3, the rate of LALA oxidation increased throughout the 40-min time course by a log_2_-fold increase of six, while the uptake rate of LALA remained relatively constant throughout, between 12 – 17 μmol C/gDW/min (Figure 2C-D, Figure S1). To confirm that differences in uptake and oxidation patterns between the strains were due to exposure to LALA and not some other experimental artifact, we measured uptake and oxidation of 4 μM [^14^C]LALA using cells pre-conditioned to 4 μM of LALA. In preconditioned HTCC7211 cells, uptake and oxidation rates remained constant, between 0.5 – 0.6 and 0.3 – 0.5 μmol C/gDW/min for uptake and oxidation rates, respectively (Figure 2I-J). In pre-conditioned HOT1A3 cells, both oxidation and transport rates stayed relatively constant after 1 min at 12 and 7 μmol C/gDW/min for uptake and oxidation rates, respectively (Figure 2G-H). Most strikingly, in pre-conditioned HOT1A3 cells, 71% of LALA had been oxidized after 5 minutes, compared to 21% at 5 min in naïve cells, indicating that pre-exposure increased the oxidative capacity of HOT1A3 cells to levels that naïve cells took 40 minutes to reach (Figure 2G-H).

We used reverse transcription quantitative real-time PCR (RT-qPCR) to confirm that transcription of the alanine dehydrogenase gene (*ald*) increased in response to LALA addition in HOT1A3 and not in HTCC7211, and to examine the timing of transcript production (Figure 3). In HOT1A3, transcripts for *ald* increased immediately upon addition of 4 μM LALA, with the transcription rate peaking at 2.5 min and remaining constant afterwards (Figure 3A). In HTCC7211, transcripts were only measured at 0 and 5 min as the phenotype data did not support an increase in *ald* transcript levels. No difference was observed between the experimental (4 μM LALA added) and control groups (same volume of water added) at 5 minutes, indicating constitutive expression of *ald* (Figure 3B).

**Figure 3.**
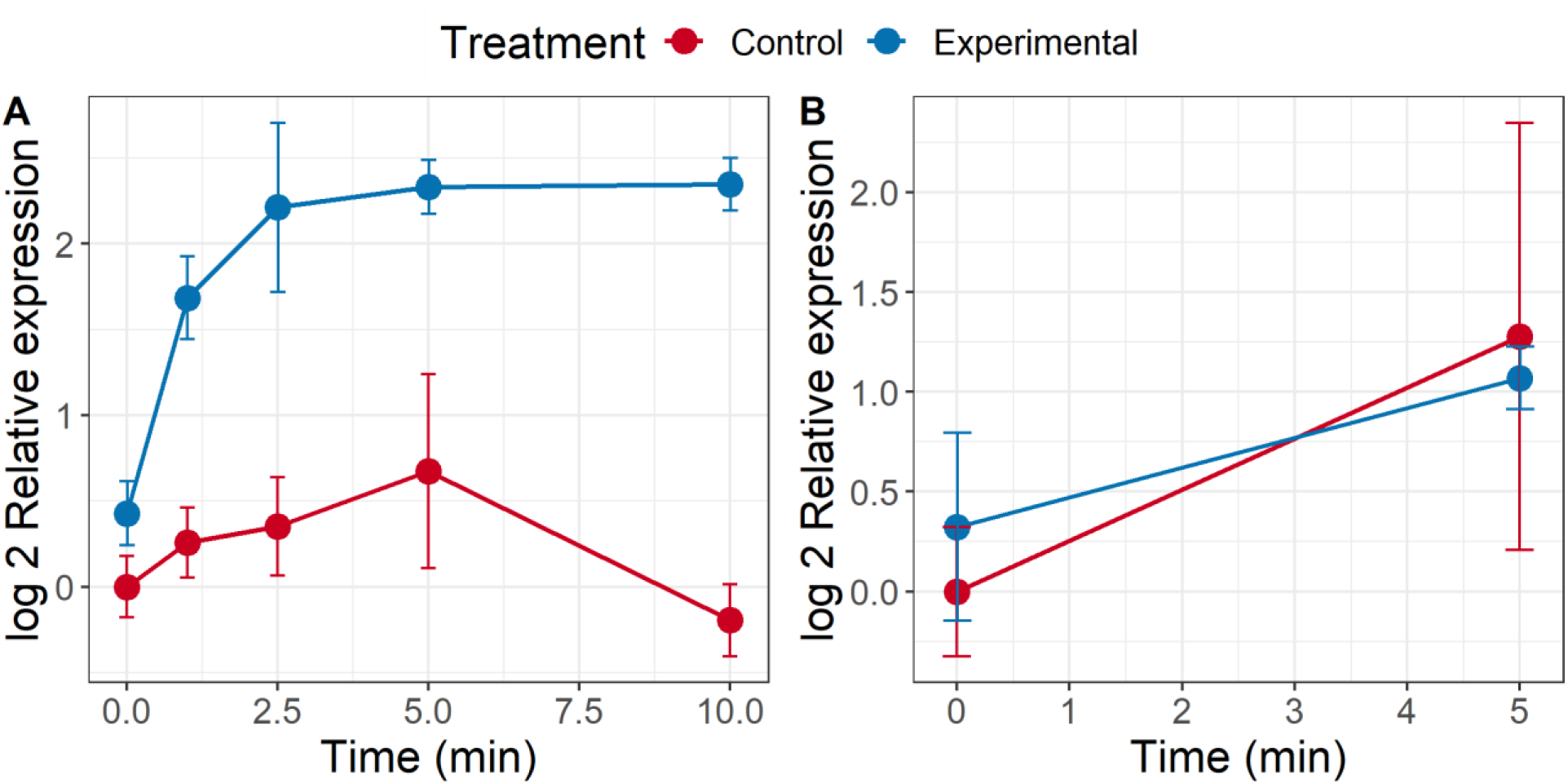
Results from a reverse transcription quantitative real-time PCR (RT-qPCR) experiment measuring transcripts for the alanine dehydrogenase gene (*ald*) in (A) *A. macleodii* st. HOT1A3 or (B) *Cand*. P. st. HTCC7211. Experimental cultures in both had 4 μM L-alanine added; an equal volume of water was added to the control cultures. *ald* transcript abundances were normalized to the abundances of two endogenous control genes, *recA* and *rpoD*. Final abundances were further normalized to the average of the control time zero abundances. Error bars are the standard deviation of four replicates: in HOT1A3, duplicate biological replicates were each split into duplicate technical replicates; in HTCC7211, quadruplicate biological replicates were used for each time point.

Based on these results, we conclude that oxidation of LALA in HOT1A3 is under transcriptional regulation, as indicated by the sharp and continual increase in oxidation rates after 5 min (Figure 2C-D), the consistent oxidation rate in pre-exposed cells (Figure 2G-H, Figure S1), the increase in *ald* transcripts upon LALA addition (Figure 3A), and the presence of a gene in the HOT1A3 genome that is homologous to the *aldR* regulator of L-alanine metabolism. HTCC7211 does not have transcriptional regulation of LALA metabolism, as indicated by the steady rate of oxidation upon LALA exposure (Figure 2A-B), the similarity in oxidation rates between naïve and pre-exposed cells (Figure 2B, F; Figure S1), the constant expression of *ald* transcripts (Figure 3B), and the absence of a homolog for the *aldR* regulator in the HTCC7211 genome.

### Experimental Observations of L-alanine Uptake and Metabolism: Measurements of Timing

Having confirmed the presence/absence of our trait of interest, we next measured the time it took both strains to initiate oxidation of LALA at a variety of LALA concentrations. Initiation of oxidation was defined as the first time point when the amount of [^14^C]LALA oxidized was greater than zero and the standard deviation did not overlap zero (marked by * in Figure 2E-L and Figure 4A-B).

**Figure 4.**
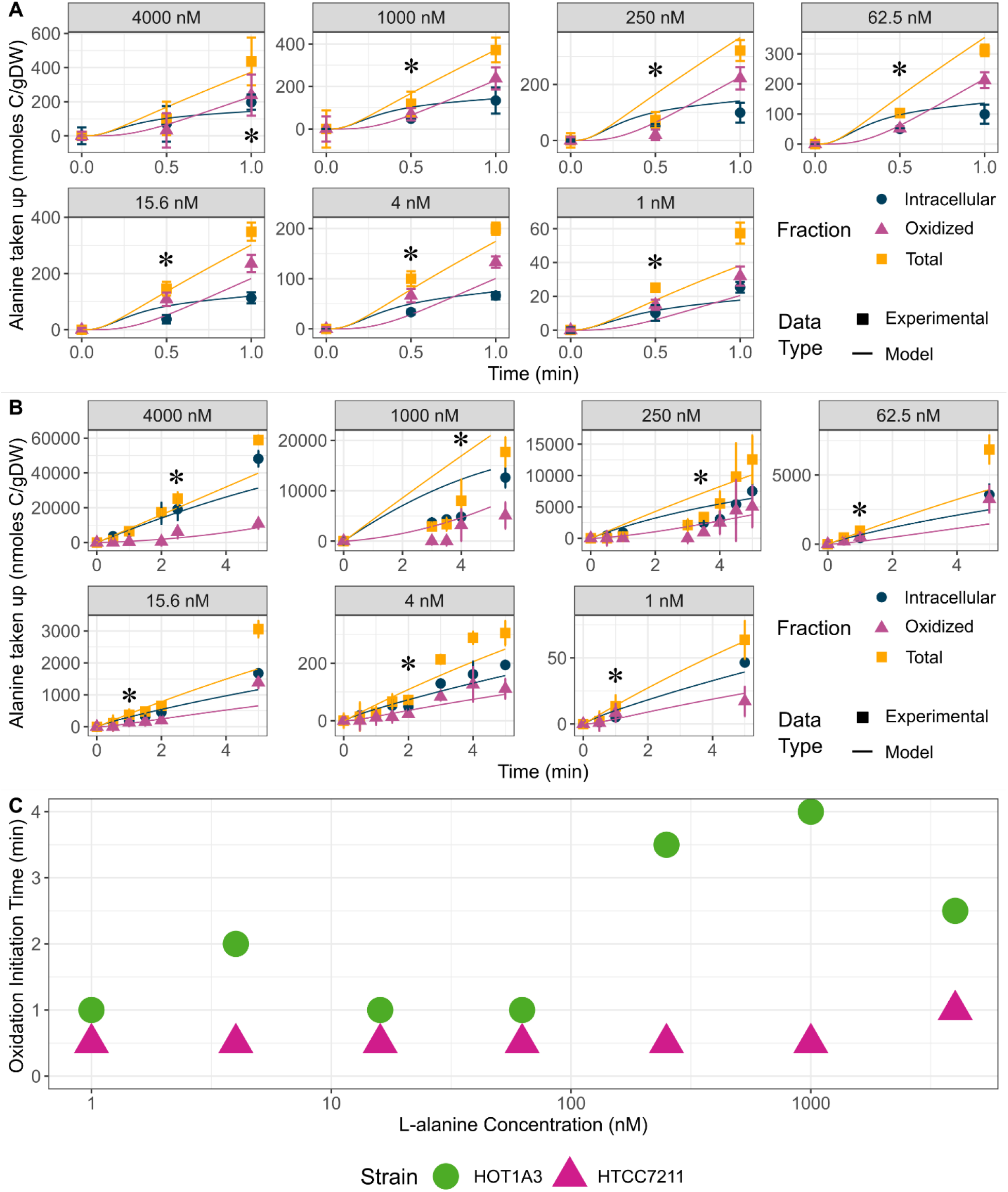
Uptake and oxidation of [^14^C]L-alanine ([^14^C]LALA) in (A) an oligotroph, *Ca. P*. st. HTCC7211, and (B) a copiotroph, *A. macleodii* st. HOT1A3, to determine initiation of oxidation at various concentrations. Points are experimental data, while lines are the output of a mechanistic model of intracellular LALA accumulation and metabolism in these strains. Cells used were naïve to LALA prior to harvesting, washing, and starving. For detailed explanation of the different fractions, see Methods. Error bars on the points are the standard deviation of triplicate cultures. * indicates oxidation initiation, the first time point where oxidation was greater than zero and the standard deviation did not overlap zero. Note that the y- and x-axis scales differ between plots. (C) Plot showing average times of initiation of LALA oxidation in the two strains over the range of measured LALA concentrations, based on data presented in (A) and (B). The x-axis is in a log10 scale to allow differentiation of concentrations. No standard deviations are shown.

In HTCC7211, oxidation was found to begin between 0 and 0.5 min for all concentrations except 4 μM, where oxidation initiation was 0.5 – 1 min (Figure 2I, Figure 4A, C). We observed a small but noticeable increase in uptake and oxidation rates after 0.5 min at all concentrations above 15.6 nM (Figure 2J, Figure 4A). This increase was not observed in pre-induced HTCC7211 cells (Figure 2E-F), which implies it may be due to a regulatory mechanism. In HOT1A3 at concentrations above 62.5 nM, oxidation initiated between 2.5 – 4 min (Figure 2K-L, Figure 4B, C), while at concentrations of 62.5 nM and below, oxidation started within the first minute, except at 4 nM, where oxidation initiation was at 2 min (Figure 4B, C). We explored this shift in oxidation initiation times by repeating the experiment with 62.5 nM out to 40 min to see if there was a later time point where the oxidation rate increased (Figure 2M-N). However, oxidation and uptake rates stayed constant over the first 10 min at 1 μmol C/gDW/min for uptake and 0.5 μmol C/gDW/min for oxidation before slowing and stopping after 20 min as the [^14^C]LALA was removed from the media. The lack of increase in oxidation rate (Figure 2N) indicates transcriptional induction did not occur in HOT1A3 at 62.5 nM LALA, which is similar to ambient concentrations of LALA in the ocean (Pocklington, 1971; Lee and Bada, 1977; Mopper and Lindroth, 1982; Lu *et al*., 2014).

### Quantitative Analysis of Experimental Results

We developed a mechanistic model for regulation of uptake and metabolism of LALA in the two strains of bacteria (1) to explore whether the observed increase in uptake and oxidation rates in HTCC7211 after 30 s was due to a regulatory mechanism; and (2) to explore the lack of induction in HOT1A3 cells at LALA concentrations below 62.5 nM. This proposed model is depicted in Figure 5 and described in Supplementary Note 1; the equations and parameters are given in Tables S1-2 and the model fits to the data are shown in Figure 2 and Figure 4 (depicted by lines).

**Figure 5.**
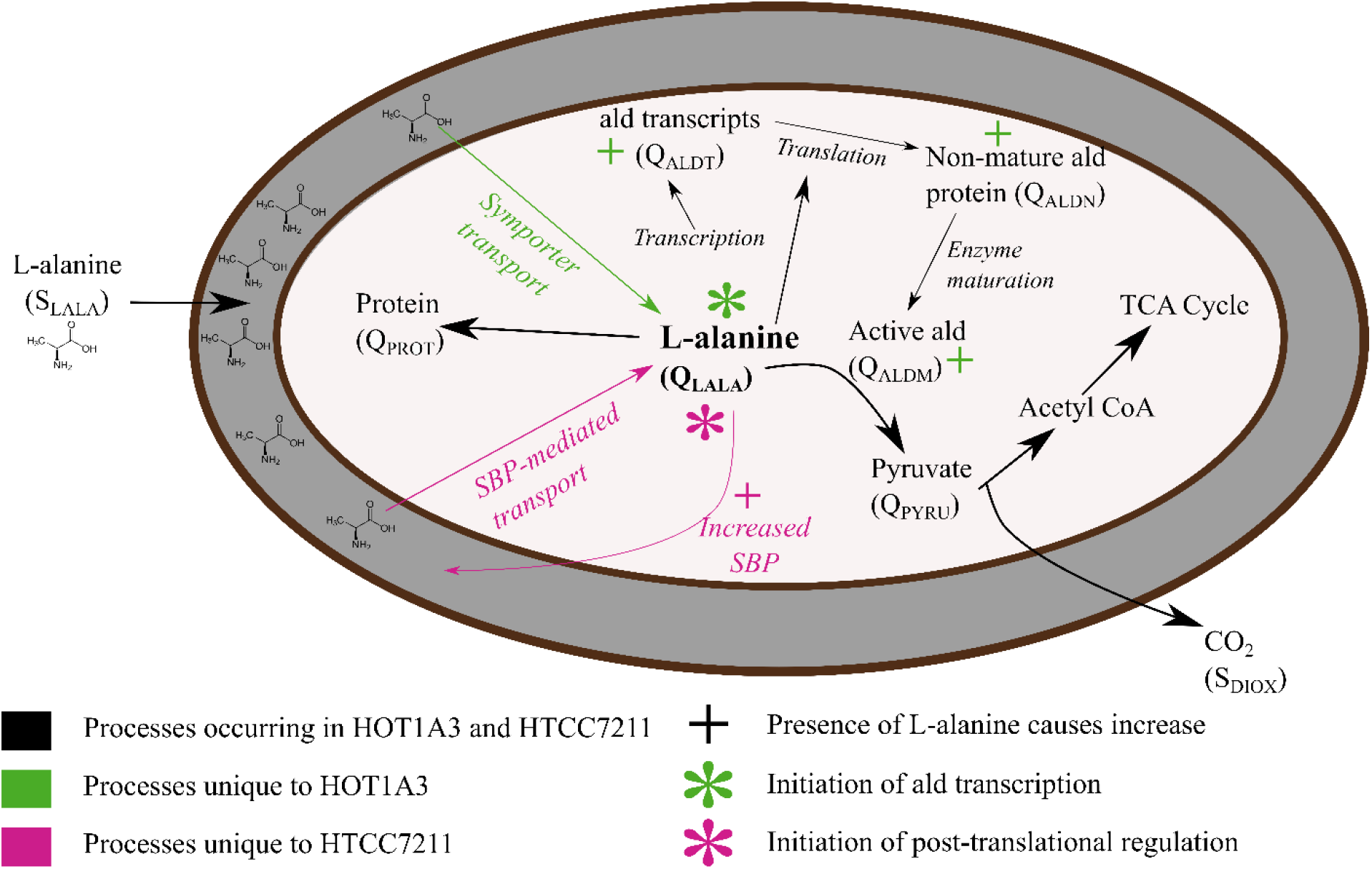
Depiction of proposed model for uptake and metabolism of L-alanine in either a copiotroph, *Alteromonas macleodii* st. HOT1A3 (green) or an oligotroph, *Ca. P*. st. HTCC7211 (pink). Transcription, translation, and enzyme maturation of alanine dehydrogenase gene (*ald*) happen constantly in HTCC7211. SBP: substrate-binding protein. For a full explanation of the model and rationale, see the text.

For (1), we hypothesized that, in HTCC7211, the experimentally observed increases in uptake rates are due to a post-transcriptional regulatory mechanism, as HTCC7211 cells preexposed to LALA exhibited no increase in transport or oxidation rates upon addition of [^14^C]LALA (Figure 2E-F, Figure S1). The increased transport of LALA would result in the increased oxidation rates observed as more LALA flows into the cell. To test our hypothesis, we optimized the fit of the model to the uptake/oxidation data either with or without regulation included in the parameters (see Methods for full description). Our model supported our hypothesis of post-transcriptional regulation in HTCC7211, since the model fit the data better with regulation included (normalized sum of error squared (SES) values: 2.53 vs. 3.78, with regulation vs. no regulation). The nature of this regulation cannot be determined from these data, but is likely post-translational, given the lengthy time (1-3 mins) required for translation and enzyme maturation (Kepes, 1963). It could be due to a post-translational modification (PTM) (Macek *et al*., 2019), a change in translational efficiency (Taylor et al., 2013; Al-Bassam et al., 2018), or an allosteric modification (Sanwal, 1970). Given the previously observed uncoupling in SAR11 between mRNA and protein levels for the LALA transporter substrate-binding protein, YhdW (Smith et al., 2013), it is likely that the regulatory mechanism increases the abundance of functional YhdW protein in response to LALA. If it is due to a PTM, N^α^-acetylation (Liang et al., 2011) is the most likely possibility, especially since there are multiple uncharacterized acetyltransferases in the HTCC7211 genome. Protein turnover rates on the order of minutes have been previously observed in other bacteria, making this mechanism plausible (Maier et al., 2011).

For (2), the mechanistic model suggested that the lack of induction in HOT1A3 cells at LALA concentrations below 62.5 nM is due to the amount of LALA entering cells not being large enough to induce transcription of ald (Figure S2). In the environment, HOT1A3 cells would not benefit from induction at ambient LALA concentrations, since they would not be bringing in enough LALA to support cellular growth. The LALA oxidation that was observed at 62.5 nM LALA and below is predicted by the model to be due to background Ald levels in the cell. This is reasonable, given that measurements of background Ald activity in *Bacillus licheniformis* (McCowen and Phibbs, 1974) would result in expected oxidation rates in HOT1A3 between 2.84 – 10.33 μmoles C/gDW/min, which exceeds the measured oxidation rate observed at 62.5 nM (0.4 μmoles C/gDW/min). Another possible explanation is that one of the two other pathways capable of metabolizing LALA in HOT1A3, via an alanine racemase or an alanineg-lyoxylate transaminase, is responsible for the oxidation measured at 62.5 nM. Follow-up transcriptome and proteome studies would be needed to evaluate the roles these pathways play.

Our model also predicts that the 2.5 – 4 min lag between addition of LALA and the initiation of LALA oxidation in HOT1A3 is primarily due to the time required for translation and protein maturation for Ald, in line with previous reports of *ald* transport and transcription taking <0.5 min (Freese and Oosterwyk, 1963; Siranosian *et al*., 1993) and translation and enzyme maturation taking at least 3 min (Pardee and Prestidge, 1961; Kepes, 1963).

### Regulation and Motility

Our primary hypothesis in this project is that the lack of transcriptional regulation in oligotrophs is related to a lack of motility/chemotaxis, which prevents oligotrophic cells from finding patches of nutrients or particles and inhabiting/attaching long enough for transcriptionally regulated systems to turn on. To test our hypothesis, we predicted residence times of motile/chemotactic (copiotrophic) and non-motile/chemotactic (oligotrophic) cells as individuals in an environment with either a continuous or instantaneous nutrient source and compared those times to our measured times for these cell types to initiate oxidation. A continuous source of nutrients would be akin to proximity to a leaking phytoplankton cell or a detritus particle, whereas an instantaneous patch would be similar to a bursting cell. Non-motile cells drifted randomly throughout the environment, simulating diffusion, while motile cells moved upgradient using a run-reverse-flick chemotaxis, which was implemented based on (Son *et al*., 2016). We present results for a range of initial distances of the bacteria to the source (Near, Middle, Far) in Figure 6B-C. A representative model run with a continuous source and cells placed a Middle distance from the source is shown in Figure 6A.

**Figure 6.**
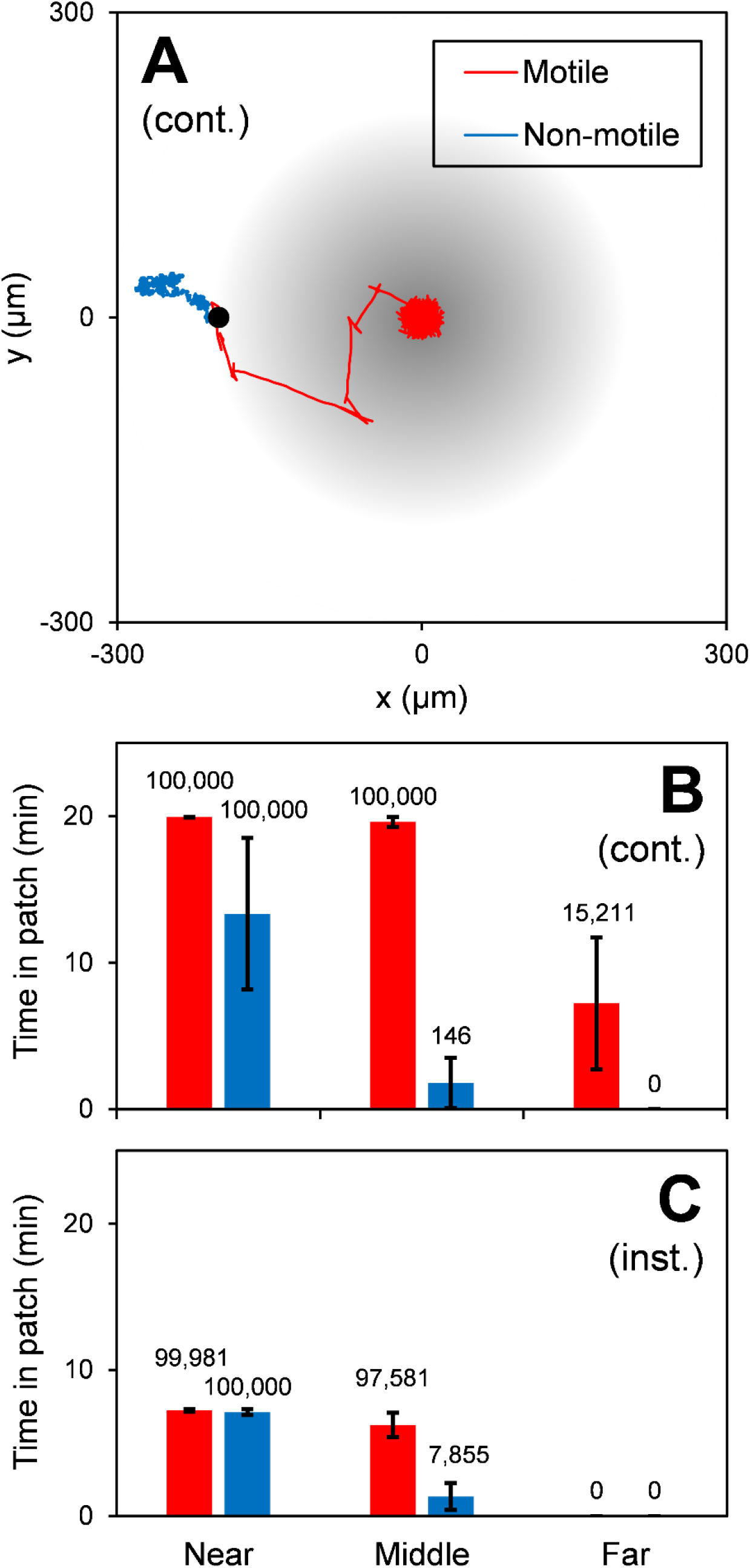
Exposure time of motile and non-motile cells to a continuous source (i.e., leaking phytoplankton cell) and an instantaneous source (i.e., lysing phytoplankton cell) of nutrients. (A) Example of cell trajectories for the continuous source scenario. Cells were initially placed 300 μm from the source (black symbol; Middle scenario). (B-C) Average time spent in patch for those cells that enter it. Cells were initially placed at 30 μm (“Near”), 300 μm (“Middle”) and 3,000 μm (“Far”) from the source. Labels indicate number of cells out of 100,000 that reach the patch. Simulations were run for 20 min., which constitutes an upper bound on the patch residence times. Error bars are standard deviation.

For the scenarios where cells are placed near the source, the cells either start in the patch (continuous source) or are quickly engulfed by the diffusing concentration (instantaneous source) (Figure 6B-C). Both cell types remain in the patch for relatively long times (>10 min). For the non-motile/chemotactic cells, this can be attributed to their lower effective diffusivity. The motile cells have a higher effective diffusivity, but that is successfully counteracted by the chemotaxis. However, the near-source scenario is unlikely and it is much more probable that the cells are further away from the nutrient source (e.g., the probability increases with the square of the distance) (Hulburt, 1970; Siegel, 1998). For the middle scenario, a larger fraction of the motile cells reach the patch than non-motile/chemotactic cells, which is probably the main benefit of chemotaxis. However, chemotaxis also affects the patch residence time, which is important for regulation. Specifically, the model predicts that, of the cells that reach the patch, the motile/chemotactic cells spend more time in the patch (>8 min) than the non-motile/chemotactic cells (<2 min). For the far distance scenario, the number of non-motile/chemotactic cells that reach the patch and the patch residence times decrease further.

The results of our simulation depend on the choice of parameters (e.g., the size of the continuous nutrient source, see Figure S6) and further modeling work will be required to fully characterize the distribution of patch residence times. However, across the tested sets of parameter values, the motile/chemotactic cell always (except the instantaneous far scenario, Figure 5C) stays in the patch long enough for cells to be fully induced and begin acquiring energy from LALA (oxidation initiation), which we measured at 2.5 – 4 min in HOT1A3. In most scenarios, the non-motile cell does not occupy the nutrient patch for more than 2 min (except the near scenarios, Figure 5B-C, and the high source size scenario, Figure S6).

The low occupancy time predicted from our model for non-motile cells supports our hypothesis that non-motile oligotrophs do not occupy nutrient patches long enough for transcriptional regulation to be viable. Using our mechanistic model, we tested what might happen if HTCC7211 cells did have transcriptional regulation of LALA metabolism (alternate parameters given in Table S2) and encountered a patch of LALA but then drifted out after 2 min (the predicted residence time of non-motile cells in a nutrient patch). We found that, with transcriptional regulation, HTCC7211 cells would experience a log_2_ fold decrease >5 in the amount of pyruvate, CO_2_, and ATP produced by cells over the course of a 40-minute simulation compared to cells without transcriptional regulation (Table S3). Using ATP levels of HTCC7211 cells under nutrient limitation (about 2.5E-7 moles ATP/gDW; (Noell and Giovannoni, 2019)), cells with transcriptional regulation would only produce 12% of their ATP stock from their encounter with the LALA patch, while cells without transcriptional regulation would produce 880% of their ATP stock. Thus, the lack of transcriptional regulation in non-motile oligotrophs like HTCC7211 and the SAR11 clade results in them not missing the opportunities they would have missed if they used transcriptional regulation.

In previous studies, clusters of chemotactic bacteria around nutrient patches were found to reach peak density within 5 min (Blackburn *et al*., 1998; Luchsinger *et al*., 1999; Stocker *et al*., 2008; Seymour *et al*., 2009), with instantaneous nutrient patches being exhausted or dispersed by shear within 10 min (Blackburn *et al*., 1998). In lab settings with limited shear, the mean residence time of individual chemotactic cells in a 0.2 mM patch of tryptone was 35 min (Yawata *et al*., 2020). Sinking particles could provide opportunities for chemotactic cells to stay at high nutrient concentrations for longer time scales. Previous studies have shown that non-motile cells have fewer encounters with nutrients, with non-motile cells encountering half (Blackburn *et al*., 1998), one-fourth (Stocker *et al*., 2008), or one-fifth (Brumley *et al*., 2019) the amount of nutrients in a patch as motile cells. We note that not all microbes that are capable of motility express their motility genes continuously (e.g., Roseobacters (Sule and Belas, 2013)) due to energetic costs, a factor that we did not account for in our model. One study found that only ~10% of a marine microbial community is motile, but this number increases dramatically when nutrient concentrations increase (Mitchell *et al*., 1995).

The absence of the motility trait in many oligotrophs, such as SAR11, may be linked to small cell size. The small cell size of SAR11 cells is proposed to be part of a suite of adaptations that confer favorable transport kinetic properties, including reduction of diffusion limitation and maximization of surface area-to-volume ratio (Schulz and Jørgensen, 2001; Andersen *et al*., 2016), thereby maximizing their fitness in competition for dissolved nutrients at the lower end of the concentration frequency spectrum (Giovannoni, 2017; Zhao *et al*., 2017). Additionally, it has been suggested that motility would be ineffective in very small cells like SAR11 because of Brownian effects (Mitchell, 1991; Dusenbery, 1997). While microenvironments and horizontal gene transfer conspire to assure a wide assortment of trait combinations will be found, our interpretation is that the exceptional prevalence of SAR11 is evidence that the niche of competing for nutrients at ambient background concentrations is relatively large in comparison to the alternatives, which typically involve finding favorable patches. However, some oligotrophic bacteria, for example some members of the SAR92 clade, are motile. SAR92 was reported to have minimal transcriptional regulation (Cottrell and Kirchman, 2016), but those experiments reported variation in transcription between cells in exponential and stationary growth phases, rather than responses to the availability of carbon resources. In *Pelagibacter*, a similarly small level of transcriptional (Cottrell and Kirchman, 2016) and proteomic (Sowell *et al*., 2008) changes were observed upon entering stationary phase, which is attributable to the lack of σ^S^ (*rpoS* gene), the stationary phase sigma factor found in many proteobacteria (Sowell *et al*., 2008). If the relationship between motility and regulation that we describe is generally true, then it would be expected that a motile oligotroph like SAR92 would have regulation of carbon assimilation pathways but would lack σ^S^, like *Pelagibacter*. In the case of the L-alanine dehydrogenase regulator, *aldR*, many SAR92 genomes in the NCBI database are predicted to harbor this gene (amino acid cover > 80%, identity >40%), while only half of these SAR92 genomes contain a homolog for *rpoS*, supporting this hypothesis.

## Conclusion

With experiments and models, we demonstrate that, as “feast and famine” bacteria, copiotrophs can employ motility to stay in patches of nutrients long enough to sense compounds and increase metabolic activity using transcriptionally regulated systems. Between patches or particles, they leave many catabolic systems turned off, minimizing the cost of synthesizing un-needed proteins. Oligotrophs, as “fast and famine” bacteria, are at the whim of their environment and experience random encounters with nutrient patches they briefly exploit before drifting out of the patch. We show that these cells constitutively express genes involved in carbon uptake and metabolism, not missing fluctuating opportunities to transport and metabolize the nutrients they need to survive, but likely failing to make the most of unusual episodes of high nutrient concentrations. Notwithstanding their limited transcriptional responses, oligotrophs may have post-transcriptional, and in this study, possibly post-translational, controls on metabolism that enable them to respond more rapidly but with much smaller amplitude to fleeting opportunities, as observed before (Meyer *et al*., 2009; Scanlan *et al*., 2009; Tripp *et al*., 2009; Ottesen *et al*., 2014; Cottrell and Kirchman, 2016; Lankiewicz *et al*., 2016; Held *et al*., 2019), in accordance with predictions (Button, 2004). Our findings add to conceptual models by linking the non-motile, small genome, solitary cell phenotype often observed in planktonic oligotrophs to a strategy of continuously expressing genes for carbon oxidation pathways.

## Methods

### Bioinformatics

To compare the prevalence of motility and chemotaxis genes in oligotrophs and copiotrophs, metagenome assembled genomes (MAGs) from the Tully et al. marine data set (Tully *et al*., 2018) were downloaded from the NCBI assembly database using get_assemblies v 0.8.6 using the settings ‘assembly_ids --function genomes -o fna --force’ (https://github.com/davised/get_assemblies). This set of MAGs was selected as it represents a broad range of marine habitats, as opposed to, e.g., the single-cell genomic data set produced from marine tropical samples only (Pachiadaki *et al*., 2019). Although MAGs can suffer from contamination and incompleteness issues, at a broad functional level, MAGs generally provide similar results to SAGs (Alneberg *et al*., 2018). One genome was randomly picked from each genus, for a total of 250 genomes (for a list of genomes, see Table S4). Proteins were annotated using bakta v 1.4.0 with default parameters (Schwengers *et al*., 2021). The commandline rpsblast v 2.13.0 was used to search annotated protein sequences as queries against the latest (date 2021-02-22) COG database subset of the Cdd (Camacho *et al*., 2009; Lu *et al*., 2020). Tabular output from the rpsblast was converted from Cdd ID to COG ID using the data from the COG2020 release from NCBI (Galperin *et al*., 2021). For each protein, COG functions were assigned using a two-tier filtering approach. The assigned bitscore cutoffs for each COG in the 2020 release were used as a first step filter, where COG hits below the bitscore cutoff were discarded. Then, of the COG hits remaining, the COG with the highest bitscore was assigned to each protein.

COG Categories were assigned based on the COG2020 annotations for each COG. Genomes were then categorized as copiotrophs or oligotrophs based on their predicted maximal growth rates, which were taken from (Weissman *et al*., 2021). The proportion of genes within each category was calculated for each genome; significant enrichment was tested using Mann– Whitney tests with a Benjamini–Hochberg correction (α=0.05). Only the COG categories of relevance to our hypotheses are shown for simplicity. To compare flagellar motility capacity between copiotrophs and oligotrophs, the numbers of flagellar genes within COG category N or within the COG categories containing flhA or flgA were calculated for each genome. Significant enrichment was tested for as before, using a Mann-Whitney test with Benjamini-Hochberg correction.

### Growth and Washing of Cultures and Uptake Experiments

Cultures were grown on artificial seawater (ASW) and washed as described more fully in the Supplemental Methods (Carini *et al*., 2013). Both strains were grown in accordance with previous reported culturing conditions (Carini *et al*., 2013, 2014; Fadeev *et al*., 2016). Preexposed cultures had 4 μM LALA two generations before harvesting and washing. Cultures were harvested between mid-exponential phase and early stationary phase (see Figure S3 for representative growth curves) using centrifugation, then washed using ultracentrifugation in ASW with no added organics.

Uptake experimental methodology is described fully in Supplemental Methods. Briefly, washed cultures were pooled to the desired volume and divided into BSA-coated septum vials and sealed with PTFE-faced butyl septa (Millipore-Sigma, 27201) (Halsey *et al*., 2012). A portion of the cultures were heated at 50°C for 1 h to make the killed-cell controls. Metabolic activity of cultures was slowed by chilling at 4°C in the dark for 1 h. Cultures were warmed to room temperature (22°C) prior to the start of uptake experiments. Reagents were added to cultures via injection with syringe and needle through the stoppers. [^14^C]L-alanine (American Radiolabeled Chemicals, ARC 0231, [1-14C]) was added to the cultures and the cultures were incubated in a dark water bath at 25°C. The reaction was stopped by the addition of either 10 mM sodium azide (for the intracellular fraction) or 0.05 N NaOH, 2.5 mM Na_2_CO_3_, and 50 mM BaCl_2_ (for the total fraction: intracellular + oxidation to ^14^CO_2_). Intracellular fraction cultures were then placed on ice and filtered within 2 h, while total fraction cultures were placed at 4°C overnight before filtering (Halsey *et al*., 2012). Cultures were filtered and washed to remove extracellular ^14^C label, then immersed in scintillation fluid (Ultima Gold XR, Perkin Elmer) and left overnight prior to counting. Resulting data were plotted using the ggplot2 package in the R software environment (R Core Team; Wickham, 2016).

### RNA extraction and RT-qPCR

RNA was extracted from HOT1A3 and HTCC7211 cells to measure transcript changes in *ald* in response to LALA addition, as described fully in the Supplemental Methods. High cell density cultures of HOT1A3 and HTCC7211 were prepared as described above; 4 μM LALA was added to experimental cultures, while an equal volume of sterile water was added to negative control cultures. Uptake was stopped by placing on ice, then pelleted via centrifugation at cold temperatures. Pellets were washed in RNAlater solution (Life Technologies, AM7020), then flash frozen in liquid nitrogen and stored at −80°C until RNA extraction was performed. RNA extraction was performed using an RNeasy min-elute clean up kit (Qiagen, 74204) according to manufacturer’s guidelines. Contaminating gDNA was removed using the Thermo Fisher Turbo DNA-free kit (Life Technologies, AM1907). Total RNA was quantified via Qubit fluorimeter (Thermo Fisher) and RNA quality assessed using the BioAnalyzer 2100 (Agilent) RNA Nano Chip at the Center for Genome Research and Biocomputing Core Facilities at Oregon State University.

For reverse transcription quantitative real-time PCR (RT-qPCR), the procedure used was similar to (Schuster, 2011), with a full description in the Supplemental Methods. RNA was reverse transcribed into cDNA using the Qiagen QuantiTect RT kit (Qiagen, 205311), including negative controls. Primers were designed for the *ala* gene and two endogenous control genes (*recA* and *rpoD*) using the Geneious Primer Design tool and ordered from Integrated DNA Technologies. Primer sequences are given in Table S5. Primers were validated on DNA from the organism of choice (*A. macleodii* st. HOT1A3 or *Cand*. P. st. HTCC7211) using PCR. qPCR was performed using the Qiagen QuantiTect SYBR Green PCR kit (Qiagen, 204143) on an Applied Biosystems 7500 Real-Time PCR System. Instrumental qPCR parameters are as follows: initial activation (95°C, 15 min), 40X cycles of denaturation (94°C, 15 s), annealing (60°C, 30s), and extension (72°C, 30 s), followed by a standard melt curve analysis. No-template controls were included to assess contamination. Primer efficiencies were measured for each HOT1A3 primer and were all above 93%. Endogenous control genes were tested for constant expression (Figure S4F-G). Data analysis was conducted in Microsoft Excel using the method described by (Vandesompele *et al*., 2002) for normalizing gene expression to multiple endogenous control genes.

### Mechanistic Model

A mechanistic model of LALA uptake and metabolism, as well as regulation of those processes, was developed for the two strains of bacteria used in this study. A list of the equations used can be found in Table S1. The model was coded into Excel and parameterized using values from literature and from experiments (Table S2). The parameters were optimized to fit all experimental data using the SOLVER add-in in Excel by setting it to minimize the summated, normalized sum of error squared value (SES) from the fit of the model to all experimental data. To calculate the SES, the experimental and model values for each experiment were first converted to molC/L, then normalized using the mean of all experimental values from that experiment. The SES was calculated using these normalized values, then divided by the number of data points for that experiment, so that experiments with larger number of data points would not have oversized influence on the optimization. SOLVER was run with the following settings: Evolutionary solving method, convergence of 1E-9 (to allow for optimizing low Km values), mutation rate of 0.075, population size of 500, maximum time without improvement of 240 s, with bounds on variables and automatic scaling. Multiple runs of SOLVER were required, using different starting points, to achieve an optimal fit of the model to the data. The starting state for the model for the pre-induced experiments was set to be equal to the final state from the model fit to the 4 μM 40-min experiment.

We tested for the presence of regulation in HTCC7211 and HOT1A3 on LALA uptake and *ald* transcription, respectively. In HTCC7211, *K_a,tran,LALA_* was set very low (1E-20) and *n_a,tran,LALA_* was set high (15), so that the activation term for uptake equaled one. Similarly, in HOT1A3, *K_a,rpom,LALA_* and *n_a,rpom,LALA_* were also set so that the activation term for transcription of *ald* equaled one, and cellular levels of *ald* transcript, naïve protein, and mature enzyme (*Q_aldT(2)_, Q_aldN(2)_*, and *Q_aldM(2)_*, respectively) were set to be equal to the maximum values from the model fit to the 4 μM 40-min experiment, to reflect a state of constitutive expression.

### Estimation of Patch Residence Time

The purpose of the patch residence model is to estimate the residence time of motile and non-motile bacteria in instantaneous (i.e. a lysing phytoplankton cell) and continuous (i.e. exudation from a live phytoplankton cell) DOM patches (see Supplemental Methods for full description). Three-dimensional and time-variable (for the instantaneous source) substrate concentrations are computed. Trajectories of motile and non-motile bacteria are computed and their patch residence time is computed based on a threshold concentration (*C_in_*, molC/m^3^). Motile and non-motile bacteria are modeled as individual agents. They diffuse using a random walk (Hellweger and Bucci, 2009) based on a diffusion coefficient (*D*, m^2^/s). In addition, motile bacteria move with a run-reverse-flick mechanism, which was implemented based on (Son *et al*., 2016). A benchmark test of our model against experimental results from that paper is shown in Figure S5B and a benchmark test of the run & tumble component of our model against the model in (Jackson, 1987) is shown in Figure S5A.

Model parameters were chosen to represent a realistic scenario (Table S6, Supplemental Methods). Diffusivities of motile and non-motile bacteria were estimated from observed cell sizes for copiotrophs and oligotrophs (Table S6, Supplemental Methods). The concentration marking the boundary between inside and outside of the patch (*C_in_*, used for Figure 5B-C) was taken at 10 μmolC/L (10,000 nmolC/L), a value considerably above background DOM concentration for the open ocean (0.1 – 50 nmolC/L, (Taylor and Stocker, 2012)). The instantaneous release mass (*M*) was taken to be large enough so that the patch exists (max. *C* > *C_in_*) longer than the time it would take cells to transcriptionally upregulate their transport system.

## Data Availability

All experimental data (uptake and qPCR data) and models (biochemical and patch models) as well as the associated scripts for analysis and plotting the figures are freely available on Github (https://github.com/stenoell/Noell-oligo-copio-regulatory-strategies). The patch model is available on a separate Github page (https://github.com/fhellweger).

## Acknowledgments

The authors thank Dr. Daniel Sher at the University of Haifa for providing the *Alteromonas macleodii* st. HOT1A3 strain for this project. The authors also thank Dr. Huw Richards for his assistance with the flagellar genes comparison. This work was funded by the National Science Foundation grant IOS-1838445.

## Notes

### Competing Interest Statement

The authors have declared no competing interest.

### Summary of Updates

Main revisions: text condensed, Figure 1 updated with new analyses of differences in flagellar presence between oligotrophs and copiotrophs.

