## Supplemental information for "Differences in the regulatory strategies of marine oligotrophs and copiotrophs reflect differences in motility"

**Supporting Information**

**Supplemental Note 1**

*Description of Proposed Model for Regulation of Uptake and Metabolism of L-alanine*

In our model, in both strains, extracellular LALA is transported into cells after diffusion into the periplasmic space either via the alanine:cation symporter (HOT1A3) or the substrate-binding protein (SBP)-dependent ABC amino acid transporter, YhdWXYZ (HTCC7211). We reasoned that there is no regulation of LALA uptake in HOT1A3, as there was a similar increase in uptake rate in both naïve and pre-conditioned cells after 10 min (Figure S1). Some increase in both uptake and oxidation rate would be expected, as the cells power up after being starved. On the other hand, we hypothesized that there is regulation of LALA uptake in HTCC7211, as described in the Results.

Once inside cells, LALA is either incorporated into cellular protein during translation or metabolized to pyruvate by the alanine dehydrogenase enzyme (Ald). Based on our experimental evidence detailed in the Results, in the HTCC7211 model, transcription and translation of the alanine dehydrogenase gene (*ald*) are continuous, while in HOT1A3, transcription of *ald* starts immediately once LALA is present in the cell (Kepes, 1963; Siranosian *et al.*, 1993). As mRNA is produced, translation occurs on ribosomes, incorporating intracellular LALA for 45/377 amino acid residues in Ald. Translated alanine dehydrogenase proteins (AldN) then undergo enzyme maturation, likely consisting primarily of subunit assembly, since active Ald (AldM) is comprised of six identical subunits (Porumb *et al.*, 1987).

**Supplemental Methods**

*Growth and Washing of Cultures*

*Candidatus* Pelagibacter sp. HTCC7211 and *Alteromonas macleodii* st. HOT1A3 were both originally isolated from open ocean, oligotrophic sites (Sargasso Sea and HOT station ALOHA, respectively) (Stingl *et al.*, 2007; Sher *et al.*, 2011). Cultures were grown on artificial seawater (ASW) (Carini *et al.*, 2013). SAR11 cells were grown in ASW amended with 100 µM pyruvate, 50 µM glycine, 10 µM methionine, and SAR11-specific vitamins (Carini *et al.*, 2013, 2014). HOT1A3 cultures were grown in ASW amended with 1 mM acetate, lactate, pyruvate, 0.5 mM glycerol, and a general vitamin mix (Fadeev *et al.*, 2016). For experiments where cultures were pre-exposed to LALA prior to washing, 4 µM LALA was added to the cultures two generations before harvesting. For HTCC7211, with an average doubling time of 1.4 days, LALA was added 3 days prior to harvesting, while LALA was added 3 hours prior to harvesting HOT1A3 cultures, as their average doubling time was 1.5 h.

Cultures were grown at 25˚C in a shaking incubator at 85 RPM in 12 h light/dark to late exponential phase before harvesting. Cells were harvested to concentrate cells and remove extracellular organics using centrifugation on a Beckman J2-21 Centrifuge at 10˚C at 30,000 g for either 90 min (HTCC7211) or 30 min (HOT1A3), followed by washing in ASW with no added organics in a Beckman-Coulter Ultracentrifuge at 10˚C at either 48,000 g for 60 minutes (HTCC7211) or 28,000 g for 20 min (Noell and Giovannoni, 2019). Supernatant was removed and the cells were re-suspended in ASW with no added organics. The centrifugation protocol used here matches that used previously in our lab, where cell viability was tested using flow cytometry and growth experiments (Giovannoni *et al.*, 2019; Noell and Giovannoni, 2019; Noell *et al.*, 2021). In previous tests, no changes in forward and side scatter or DNA fluorescence peaks were observed using flow cytometry in SAR11 cells that underwent these treatments. SAR11 cells that underwent these treatments grew normally when put back into growth media (see the positive control cultures in the growth experiments in (Noell and Giovannoni, 2019)).

*Uptake Experiments*

Washed cultures were pooled to the desired volume at a cell density between 2 – 8 E7 cells/mL and cell counts taken, as previously described (Carini *et al.*, 2013). Pooled cultures were divided into appropriate volumes in individual BSA-coated septum vials and sealed with PTFE-faced butyl septa (Millipore-Sigma, 27201) (Halsey *et al.*, 2012). A portion of the cultures were heated at 50˚C for 1 h to make the killed-cell controls. Metabolic activity of cultures was slowed by chilling at 4˚C in the dark until counting finished, 1 h, to induce a partial starvation state similar to that experienced by most cells in the ocean (Morita, 1982; Moriarty and Bell, 1993). Cultures were kept in the dark throughout to remove any possibility of cells gaining energy from rhodopsins (Steindler *et al.*, 2011). Cultures were warmed to room temperature (22℃) using a heater prior to the start of uptake experiments. As above, these steps follow previously used protocols and the impact of cooling/heating on cell viability was minimal (Noell and Giovannoni, 2019). Reagents were added to cultures via injection with syringe and needle through the stoppers. [$^{14}$C]L-alanine (American Radiolabeled Chemicals, ARC 0231, [1-14C]) was added to the cultures at the desired final concentration. Cultures were then incubated for the needed time in a dark water bath at 25˚C. The reaction was stopped by the addition of either 10 mM sodium azide (for the intracellular fraction) or 0.05 N NaOH, 2.5 mM $Na_2CO_3$, and 50 mM $BaCl_2$ (for the total fraction: intracellular + oxidation to $^{14}CO_2$). Intracellular fraction cultures were then placed on ice and filtered within 2 h, while total fraction cultures were placed at 4˚C overnight before filtering (Halsey *et al.*, 2012). Cultures were filtered on 0.1 or 0.2 µm PTFE Omnipore Membrane filters (HTCC7211 and HOT1A3, respectively), washed three times with 3 mL of un-amended ASW to remove extracellular $^{14}$C label, and transferred to scintillation vials with scintillation fluid (Ultima Gold XR, Perkin Elmer) and left overnight prior to counting.

Triplicate cultures were used for each time point for each fraction, for a total of six cultures for each time point. Raw counts from a blank filter were subtracted from each sample. To calculate the $^{14}CO_2$ produced (oxidized fraction), the $^{14}$C signal in the intracellular fraction was subtracted from the total fraction. Average $^{14}$C signal in the time-zero time points for each fraction was subtracted from all other time points in that fraction to account for variability between the two methods used, as it takes longer for the azide reagent to be added than the multiple reagents required for the $BaCl_2$ method. Total uptake rates were calculated by dividing differences in total fraction values by difference in the time between those points. The total fraction was used for this as it represents the total amount of [$^{14}$C]LALA that fluxed into the cell. Oxidation rates were calculated in the same way using the oxidation values for time points. Figures were plotted using the ggplot2 package in the R software environment (Wickham, 2016; R Core Team).

*RNA extraction and RT-qPCR*

RNA was extracted from HOT1A3 and HTCC7211 cells to measure transcript changes in *ald* in response to LALA addition. High cell density cultures of HOT1A3 and HTCC7211 were prepared as described above, final volume of 10 mL (final cell density of ~1E10 cells/mL). To the experimental cultures, LALA was added at 4 µM, while an equal volume of sterile water was added to negative control cultures. Cultures were incubated as described above for various times, then placed on ice to stop cell activity. Cells were then pelleted via centrifugation, maintaining 0˚C temperature throughout. Pellets were re-suspended in RNAlater solution (Life Technologies, AM7020) at room temperature for 15 min, pelleted again via ultracentrifugation, then flash frozen in liquid nitrogen following supernatant removal and stored at -80˚C until RNA extraction was performed. For HOT1A3, biological duplicates were performed for each time point and treatment group, which were each split into technical duplicates prior to RNA extraction, for a total of 4 replicates. For HTCC7211, biological quadruplicates were performed for the two time points.

RNA extraction was performed using an RNeasy min-elute clean up kit (Qiagen, 74204) with an on-column DNase treatment (Qiagen, 79254) and a 5 minute incubation with buffer RW1 to enhance genomic DNA (gDNA) removal per the manufacturer's instructions. Samples were eluted in 20 µL of RNase-free water and stored at -80˚C until further analysis. Remaining contaminating gDNA was removed using the Thermo Fisher Turbo DNA-free kit (Life Technologies, AM1907), following the manufacturer's recommendation of carrying out two separate incubations with half the total DNase (30 min each) on each sample. Total RNA was quantified via Qubit fluorimeter (Thermo Fisher) and RNA quality assessed using the BioAnalyzer 2100 (Agilent) RNA Nano Chip at the Center for Genome Research and Biocomputing Core Facilities at Oregon State University. All samples had no signs of RNA degradation and had rRNA ratios between 1.5 and 2.

For reverse transcription quantitative real-time PCR (RT-qPCR), the procedure used was similar to (Schuster, 2011). 300 ng of RNA from each sample was reverse transcribed into cDNA using the Qiagen QuantiTect RT kit (Qiagen, 205311) following the kit instructions, final volume of 20 µL. Negative controls with no template RNA added and with no reverse transcriptase added were included to check for contaminating gDNA or procedural contamination, which was checked with PCR and gels; no contamination was observed (data not shown). Primers were designed for the *ala* gene and two endogenous control genes (*recA* and *rpoD*) with $T_m$ values close to 65˚C using the Geneious Primer Design tool, which screens for off-target sites. Primers were ordered from Integrated DNA Technologies with standard desalting. Primer sequences are given in Table S5. Primers were validated on DNA from the organism of choice (*A. macleodii* st. HOT1A3 or *Cand.* P. st. HTCC7211) using PCR; no off-target amplification was observed, with a single, strong band for each primer in each strain (data not shown). qPCR was performed using the Qiagen QuantiTect SYBR Green PCR kit (Qiagen, 204143) on an Applied Biosystems 7500 Real-Time PCR System, with a final reaction volume of 25 µL; all reagents followed the manufacturer's recommendations. A primer concentration of 0.2 µM was used and 1 µL of 1:10 diluted cDNA sample was added. Optical 96-well reaction plates (Life Technologies, 4346906) were sealed with clear adhesive film (Life Technologies, 4306311) and centrifuged briefly prior to placing into instrument. Instrumental qPCR parameters are as follows: initial activation (95˚C, 15 min), 40X cycles of denaturation (94˚C, 15 s), annealing (60˚C, 30s), and extension (72˚C, 30 s), followed by a standard melt curve analysis. No-template controls were included to assess contamination; in all controls, either no amplification or $C_q$ values above 37 were observed. Melt curves all showed a single peak for each primer (Figure S4A-E). Each experimental sample was measured in duplicate for qPCR. Primer efficiencies were measured for each HOT1A3 primer using a 5X 1:10 dilution series (in duplicate) of cDNA sample and were all above 93%. Endogenous control genes were tested for constant expression across all time points and experimental groups (Figure S4F-G). Data analysis was conducted in Microsoft Excel using the method described by (Vandesompele *et al.*, 2002) for normalizing gene expression to multiple endogenous control genes. The average value of the control group time zero replicates was used as the calibrator for calculating $\Delta C_T$ values. Data was extracted from runs using the 7500 FAST System software (v 2.0.6).

*Patch Residence Model: Microbial Diffusion*

Microbes diffuse using a stochastic process (Hellweger and Bucci, 2009). For example, the change in *x* (m) position in a time step (*Δt*, d) is:

$$\Delta x = r\sqrt{2\,D\,\Delta t} \tag{1}$$

where *r* is a random number drawn from a standard normal distribution and *D* ($m^2/s$) is the diffusion coefficient.

*Patch Residence Model: Chemotaxis*

The model used in the simulations uses run-reverse-flick chemotaxis, which is based on run & tumble chemotaxis. Here both submodels are described.

*Patch Residence Model: Run & tumble model*

The run & tumble chemotaxis submodel was adopted from (Jackson, 1987). See Figure S5A for a reproduction of one of the experiments in that reference. Bacteria move (i.e. run) at a specified constant velocity ($v_{ave}$, m/s) in a direction defined by their orientation ($\theta$, $\varphi$, rad). Note that a Cartesian coordinate system is used here. During the run, the orientation changes based on a rotational diffusion coefficient ($D_r$, rad$^2$/d) (see Eqn. (1)).

The probability of a tumble ($P_t$) in a timestep ($\Delta t$, d) is:

$$P_t = \frac{\Delta t}{\tau} \tag{2}$$

where $\tau$ (d) is the average run time, which is defined as:

$$\tau = \tau_0 \, exp\left(\alpha \left[\overline{\frac{dP_b}{dt}}\right]\right) \tag{3}$$

where $\tau_0$ (d) is the average run time in the absence of a concentration gradient, $\alpha$ (d) is a constant that defines the increase of $\tau$ with $\left[\overline{\frac{dP_b}{dt}}\right]$ (1/d), which is the running, exponentially decaying average of the rate of change of the fraction of bound surface receptors ($P_b$). That value is updated each time step based on the present value:

$$\left[\overline{\frac{dP_b}{dt}}\right]_{t+\Delta t} = \left(1 - \frac{\Delta t}{T_m}\right)\left[\overline{\frac{dP_b}{dt}}\right]_t + \frac{\Delta t}{T_m}\left[\frac{dP_b}{dt}\right] \tag{4}$$

where $T_m$ (d) is a time constant that defines the memory time scale. The rate of change of the fraction of bound surface receptors is:

$$\left[\frac{dP_b}{dt}\right] = \frac{K_D}{(K_D + C)^2}\frac{\Delta C}{\Delta t} \tag{5}$$

where $K_D$ (mol/m$^3$) is the half-saturation constant of the surface receptors and $C$ (mol/m$^3$) is the concentration. When a run stops (i.e. tumble), the bacteria orientation ($\theta$, $\varphi$, rad.) is randomized.

*Patch Residence Model: Run-reverse-flick & chemokinesis*

The run-reverse-flick and chemokinesis submodel is based on (Son *et al.*, 2016). See Fig. S4B for the reproduction of one of the experiments in that reference. The bacteria have a base speed in the absence of a chemoattractant ($v_i$, m/s), which varies randomly among the individuals. This is assigned at division by drawing from a lognormal distribution with assigned mean ($v_{move,ave}$, m/s) and standard deviation ($v_{logvar}$, m/s), truncated to avoid unrealistic values ($v_{mot,min}$, $v_{mot,max}$, m/s). The base speed is increased by a factor ($f_v$) if the substrate concentration exceeds a threshold ($C_v$, mol/m$^3$).

The average run time is a function of the history of concentration experienced (as in the run & tumble model above) as well as the velocity. The relationship with the velocity is based on an empirical relationship between the reorientation frequency (*f(v)* (1/s) in (9), 1/τ (s) here) and the velocity ($v_{mot}$, µm/s) (Eq. S5 in (Son *et al.*, 2016)):

$$\tau_v(v_{mot}) = \frac{c1}{1 + e^{c2\,(v_{mot}-c4)}} + c3 \tag{6}$$

where *c1* ($\eta$ in (9), s), *c2* ($\zeta$ in (9), s/µm), *c3* ($\theta$ in (9), s) and *c4* ($v_t$ in (9), µm/s) are constants. The combined function is:

$$\tau = \tau_0\, exp\left(\alpha \left[\frac{dP_b}{dt}\right]\right) \frac{\tau_v(v_{mot})}{\tau_v(v_{mot,ave})} \tag{7}$$

The occurrence of a reorientation event is determined based on the above equation. When that happens, a forward-moving cell will reverse direction and start a backward run. A backward-moving cell will reverse direction and start moving forward and at that time may also flick, based on an empirical relationship between the probability of flicking and the velocity (Son *et al.*, 2016):

$$P_F = c5 + \frac{c6}{1 + e^{c7\,(v_{mot}-c8)}} \tag{8}$$

where *c5*, *c6*, *c7* (s/µm) and *c8* (µm/s) are constants.

*Patch Residence Model: Source Equations*

DOM concentrations are attenuated by diffusion (i.e. no effect of bacteria consumption). The concentration due to the instantaneous source is:

$$C(x, y, z, t) = \frac{M}{(4\,\pi\,D_{DOM}\,t)^{3/2}}\ exp\left(-\frac{x^2 + y^2 + z^2}{4\,D_{DOM}\,t}\right) + C_\infty \tag{9}$$

where $C$ (molC/m$^3$) is the DOM concentration, $M$ (molC) is the patch mass and $D_{DOM}$ (m$^2$/s) is the DOM diffusion coefficient. $C_\infty$ is the background concentration, which is assumed negligible here. The steady-state concentration gradient for the continuous source is from (Jackson, 1987):

$$C(x, y, z) = \frac{\left(\frac{W}{4\,\pi\,D_{DOM}}\right)}{\sqrt{x^2 + y^2 + z^2}} + C_\infty \tag{10}$$

where $W$ (molC/d) is the source rate.

*Patch Residence Model: Parameters*

The model equations and parameterization are generally based on the observations of (Son *et al.*, 2016). However, there is a discrepancy, because the model is three-dimensional, whereas the observations are in two dimensions. That means observed velocities do not include the third component. For a random orientation, the average ratio of 3D/2D velocities is 1.6 (based on our simulations), which was applied to the velocity parameters (e.g., $v_{mot,ave}$, $c4$). Also, some flicks may appear as reversals or continuations in two dimensions. The actual frequency of reorientation and flicking are therefore also expected to be higher, but here the magnitude is more difficult to estimate as it also depends on the measurement precision and criteria for identifying these events. Corresponding parameters (e.g., $\tau_0$, $c5$) were therefore calibrated to match observations (see Figure S5B).

*Patch Residence Model: Source Parameters*

For the continuous source, the rate is based on a large phytoplankton cell with a radius of 20 µm, growing at 1/d and exuding 20%. For the instantaneous source, the mass is based on a slightly smaller phytoplankter with a cell radius of 10 µm, where 5% of the biomass is available (i.e. DOM). Calculations are as follows:

$$W = \frac{4}{3}\pi\,(20\;\mu m)^3 \frac{(100\;cm)^3}{(10^6\;\mu m)^3} 1\frac{gWW}{cm^3} 0.1\frac{gDW}{gWW} 0.4\frac{gC}{gDW}\frac{molC}{12\;gC} 1\frac{1}{d}\frac{d}{86400\;s} 0.20 = 2.6\;10^{-16}\;\frac{molC}{s} \tag{11}$$

$$M = \frac{4}{3}\pi\,(10\;\mu m)^3 \frac{(100\;cm)^3}{(10^6\;\mu m)^3} 1\frac{gWW}{cm^3} 0.1\frac{gDW}{gWW} 0.4\frac{gC}{gDW}\frac{molC}{12\;gC} 0.05 = 7.04\;10^{-13}\;molC \tag{12}$$

**Supplemental Tables**

**Table S1** Model equations for mechanistic model of L-alanine uptake/metabolism in the two strains studied. Portions in pink or green indicate regulatory processes unique to *Ca.* Pelagibacter st. HTCC7211 or *Alteromonas macleodii* st. HOT1A3, respectively.

| Equation No. | Formula | Description |
|---|---|---|
| **Mass Balances** | | |
| 1 | $\frac{dS_{LALA(i)}}{dt} = -V_{tran,LALA(i)}\ X$ | Extracellular L-alanine (LALA) [molC/L] |
| 2 | $\frac{dQ_{LALA(i)}}{dt} = +V_{tran,LALA(i)} - \boldsymbol{f_{LALA/PROT}\ V_{rptm,PROT(i)}} - \boldsymbol{f_{LALA/ALDN}\ V_{rptm,ALDN(i)}} - V_{pyru,LALA(i)} - k_{grow}\ Q_{LALA(i)}$ | Intracellular L-alanine (LALA) [molC/gDW] |
| 3 | $\frac{dQ_{PYRU(i)}}{dt} = +V_{pyru,LALA(i)} - V_{diox,PYRU(i)} - k_{grow}\ Q_{PYRU(i)}$ | Intracellular pyruvate (PYRU) [molC/gDW] |
| 4 | $\frac{dS_{DIOX(i)}}{dt} = V_{diox,PYRU(i)}\ X$ | Extracellular CO2 (DIOX) [molC/L] |
| 5 | $\frac{dQ_{ALDT}}{dt} = V_{rpom,ALDT} - k_{deca,ALDT}\ Q_{ALDT} - k_{grow}\ Q_{ALDT}$ | Intracellular ALD transcript (ALDT) [gALDT/gDW] |
| 6 | $\frac{dQ_{ALDN(i)}}{dt} = V_{rptm,ALDN(i)} - k_{matu,ALDN}\ Q_{ALDN(i)} - k_{deca,ALDN}\ Q_{ALDN(i)} - k_{grow}\ Q_{ALDN(i)}$ | Intracellular naïve ALD protein (ALDN) [gALDP/gDW] |
| 7 | $\frac{dQ_{ALDM(i)}}{dt} = +k_{matu,ALDN}\ Q_{ALDN(i)} - k_{deca,ALDM}\ Q_{ALDM(i)} - k_{grow}\ Q_{ALDM(i)}$ | Intracellular mature ALD protein (ALDM) [gALDP/gDW] |
| 8 | $\frac{dQ_{PROT(i)}}{dt} = V_{rptm,PROT(i)} - k_{grow} Q_{PROT(i)}$ | Intracellular protein (PROT) [gPROT/gDW] |
| 9 | $\frac{dQ_{WAST(i)}}{dt} = +k_{deca,ALDN} Q_{ALDN(i)} + k_{deca,ALDM} Q_{ALDM(i)} - k_{grow} Q_{WAST(i)}$ | Waste products (WAST) [gALD/gDW] |
| 10 | $\frac{dQ_{INTR(i)}}{dt} = +Q_{LALA(i)} + Q_{PYRU(i)} + Q_{ALDN(i)} f_{LALA/ALDN} + Q_{ALDM(i)} f_{LALA/ALDN} + Q_{PROT(i)} f_{LALA/PROT} + Q_{WAST(i)} f_{LALA/PROT}$ | Total Intracellular label [molC/gDW] |
| **Rates** | | |
| 11 | $V_{tran,LALA(i)} = V_{max,tran,LALA} \frac{S_{LALA(i)}}{K_{m,tran,LALA} + S_{LALA(T)}} \left(B + \frac{Q_{LALA(T)}^{n_{a,tran,LALA}}}{K_{a,tran,LALA}^{n_{a,tran,LALA}} + Q_{LALA(T)}^{n_{a,tran,LALA}}}\right)$ | L-alanine (LALA) transport [molC/gDW/d] |

| | | |
|---|---|---|
| 12 | $V_{pyru,LALA(i)} = k_{pyru,LALA} Q_{ALDM(T)} \frac{Q_{LALA(i)}}{K_{m,pyru,LALA} + Q_{LALA(T)}}$ | L-alanine (LALA) conversion to pyruvate (PYRU) [molC/gDW/d] |
| 13 | $V_{diox,PYRU(i)} = V_{max,diox,PYRU} \frac{Q_{PYRU(i)}}{K_{m,diox,PYRU} + Q_{PYRU(T)}}$ | Pyruvate (PYRU) conversion to CO2 (DIOX) [molC/gDW/d] |
| 14 | $V_{rpom,ALDT} = V_{max,rpom,ALDT} \frac{K_{i,ALDT,ALDM}{}^{n_{i,ALDT,ALDM}}}{K_{i,ALDT,ALDM}{}^{n_{i,ALDT,ALDM}} + Q_{ALDM}{}^{n_{i,ALDT,ALDM}}} \frac{Q_{LALA(T)}{}^{n_{a,rpom,LALA}}}{K_{a,rpom,LALA}{}^{n_{a,rpom,LALA}} + Q_{LALA(T)}{}^{n_{a,rpom,LALA}}}$ | ALD transcription [gALDT/gDW/d] |
| 15 | $V_{rptm,PROT(i)} = f_{PROT/DW}\ k_{synt} \frac{Q_{LALA(i)}}{K_{m,prot,LALA} + Q_{LALA(T)}}$ | Protein translation [gPROT/gDW/d] |
| 16 | $V_{rptm,ALDN(i)} = f_{ALD/DW}\ k_{synt} \frac{Q_{LALA(i)}}{K_{m,prot,LALA} + Q_{LALA(T)}} \frac{Q_{ALDT}}{K_{m,rptm,ALDT} + Q_{ALDT}}$ | ALD translation [gALDN/gDW/d] |

**Table S2** Model parameters and values for mechanistic model of L-alanine uptake/metabolism in the two strains studied. Also included are alternate parameter values used in the model runs for: *Ca.* Pelagibacter st. HTCC7211 either without regulation of transport or with both transporter regulation and transcriptional regulation of alanine dehydrogenase (similar to that in HOT1A3); and for *Alteromonas macleodii* st. HOT1A3 without transcriptional regulation of alanine dehydrogenase. When no alternate value is given, the parameter is the same in all model runs.

| Description | Parameter Symbol | Units | HTCC7211 Value With / Without Transporter Regulation | HTCC7211 Value With Transporter and Metabolic Regulation | HOT1A3 Value With / Without Regulation | Note |
|---|---|---|---|---|---|---|
| Growth rate | $k_{grow}$ | 1/d | 0 | | 0 | |
| Synthesis rate | $k_{synt}$ | 1/d | 0.73 | 0.73 | 16.5 / 32 | a |
| Cells in culture | $X$ | gDW/L | 3.3E-4 / 5.48E-4 | 3.3E-4 | 1.7E-3 / 6.16E-4 | b |
| Amount of protein per biomass | $f_{PROT/DW}$ | gPROT /gDW | 0.61 | 0.61 | 0.63 | c |
| Amount of ALD per biomass | $f_{ALD/DW}$ | gALD/ gDW | 1.52E-2 / 2.59E-2 | 5.36E-4 | 5.36E-4 / 1.4E-3 | d |
| Alanine in cell protein | $f_{LALA/PROT}$ | molC/g PROT | 4.3E-4 | 4.3E-4 | 4.3E-4 | e |
| Alanine in ALD enzyme | $f_{LALA/ALDN}$ | molC/g ALD | 4.2E-4 | 4.2E-4 | 5.36E-4 | f |
| Maximal rate of alanine transport | $V_{max,tran,LALA}$ | molC/g DW/d | 1.12E-3 / 6E-4 | 1.12E-3 | 1.34E-2 / 2.21E-2 | g |
| Half-saturation constant for alanine transport | $K_{m,tran,LALA}$ | molC/L | 8.3E-10 / 2.12E-9 | 8.3E-10 | 2.55E-7 / 6.65E-7 | h |
| Transport induction half-saturation value | $K_{a,tran,LALA}$ | molC/g DW | 2.05E-8 / →0 | 2.05E-8 | →0 | i |
| Activation constant for regulation of uptake | $n_{a,tran,LALA}$ | - | 0.73 / 15 | 0.73 | 10 / 15 | j |
| Rate of ALD enzyme activity | $k_{pyru,LALA}$ | molC/g ALDM/ d | 4.87 / 1.5 | 4.87 | 147 / 100 | k |
| Half-saturation value for ALD enzyme | $K_{m,pyru,LALA}$ | molC/g DW | 6.04E-6 / 5.01E-6 | 6.04E-6 | 6.7E-7 / 4.14E-6 | l |

| | | | | | | |
|---|---|---|---|---|---|---|
| Maximal rate of conversion of pyruvate to $CO_2$ | $V_{max,diox,PYRU}$ | molC/g DW/d | 0.21 / 0.26 | 0.21 | 4.8 / 4.5 | m |
| Half-saturation value for pyruvate dehydrogenase | $K_{m,diox,PYRU}$ | molC/g DW | 2.58E-5 / 1.58E-5 | 2.58E-5 | 6E-5 / 7.8E-5 | n |
| Maximal rate of RNA polymerase | $V_{max,rpom,ALDT}$ | gALDT /gDW/d | 7.21E-1 / 6.73E-1 | 7.21E-1 | 9.8 / 9.5 | o |
| Half-saturation value for initiation of ald transcription | $K_{a,rpom,LALA}$ | molC/g DW | →0 | 5E-7 | 5E-7 / →0 | p |
| Activation constant for regulation of ALDT | $n_{a,rpom,LALA}$ | - | 15 | 7 | 7 / 15 | q |
| Half-saturation value of alanine-tRNA ligase enzyme | $K_{m,prot,LALA}$ | molC/g DW | 1.8E-8 / 1.08E-7 | 1.8E-8 | 1.44E-6 / 7.13E-7 | r |
| Half-saturation value of ribosomes for transcripts | $K_{m,rptm,ALDT}$ | gALDT /gDW | 7.3E-7 / 8.13E-7 | 7.3E-7 | 2.7E-7 / 2.51E-7 | s |
| Rate of ALD mRNA decay | $k_{deca,ALDT}$ | 1/d | 183 / 366 | 183 | 435 / 481 | t |
| Rate of naïve ALD protein decay | $k_{deca,ALDN}$ | 1/d | 0.74 / 0.48 | 0.74 | 0.84 / 0.3 | u |
| Rate of mature ALD protein decay | $k_{deca,ALDM}$ | 1/d | 0.74 / 0.56 | 0.74 | 0.81 / 0.78 | v |
| Rate of ALD protein maturation | $k_{matu,ALDN}$ | 1/d | 1033 / 918 | 1033 | 2408 / 2233 | w |
| Limitation term for ALDT production | $K_{i,ALDT,ALDM}$ | gPROT /gDW | 2E-2 | 5E-5 | 5E-5 / 8E-5 | x |
| Limitation constant for ALDT production | $n_{i,ALDT,ALDM}$ | - | 3 / 1.28 | 10 | 10 / 3 | y |
| Basal uptake fraction | $B$ | - | 0.02 / 0 | 0.02 | - | z |
| Extracellular LALA concentration | $S_{LALA}(1)$ | molC/L | | Varies based on experiment | | |
| Intracellular LALA | $Q_{LALA}(2)$ | molC/g DW | →0 | →0 | →0 | aa |

| | | | | | | |
|---|---|---|---|---|---|---|
| concentration prior to expt | | | | | | |
| Intracellular pyruvate concentration prior to expt | $Q_{PYRU}(2)$ | molC/g DW | →0 | →0 | →0 | aa |
| Extracellular $CO_2$ concentration | $S_{DIOX}(2)$ | molC/L | 0 | 0 | 0 | |
| ALD transcript abundance | $Q_{ALDT}$ | gALDT /gDW | 3E-6 | →0 | →0 / 1E-2 | ab |
| ALD protein non-mature abundance | $Q_{ALDN}(2)$ | gALD N/gDW | 1.65E-2 | 1.71E-6 | 1.71E-6 / 2E-4 | ac |
| Mature ALD abundance | $Q_{ALDM}(2)$ | gALD M/gD W | 2.23E-2 | 7.5E-6 | 7.5E-6 / 2E-4 | ad |

[a] Normal growth rates under nutrient replete conditions are 0.46 and 11.1 1/d for HTCC7211 and HOT1A3, respectively. In experiments, since cells were starved for 1 h, cell activity likely increased to higher levels than under nutrient replete growth, thus a range of values from normal growth rates up to 3X normal was used to calibrate the model.
[b] From experiments; calibration range was based on the cell concentrations used in experiments, 2E7 to 1E8 cells/mL.
[c] From literature (Simon and Azam, 1989); calculated based on protein per cell in marine bacteria; smaller volume bacteria have lower protein/cell; HTCC7211 biomass from (Cermak *et al.*, 2017), HOT1A3 volume from (Fourquez *et al.*, 2014), converted using a formula of fg dry weight = 218 * $V^{0.86}$ (V in $\mu m^3$) for volumes of bacterial cells measured via DAPI staining (Posch *et al.*, 2001)
[d] HTCC7211 value calculated from maximum ALD activity in *Bacillus licheniformis* divided by range of activity of ALDM, $k_{PYRU,LALA}$. HOT1A3 value calculated from background ALD activity. Both values scaled by growth rate of *B. licheniformis* (28.5 1/d), a factor of 0.016 for HTCC7211 and 0.39 for HOT1A3
[e] From literature (Neidhardt *et al.*, 1990); calculated from *E. coli*, with 9.6% relative abundance of ala among proteins by weight
[f] Calculated from ald sequences; 9.4% for HTCC7211, 11.9% for HOT1A3
[g] Literature range for marine ultramicrobacterium strain RB2256 is 1.4E-3 – 2E-2 molC/d/gDW (Schut *et al.*, 1995). 6E-4 and 3.5E-2 molC/d/gDW are the calculated values from experimental data for HTCC7211 and HOT1A3, respectively, from non-linear regression, fitting the Michaelis-Menten equation to uptake data.
[h] Value from seawater communities is 1E-9 molC/L (Button and Robertson, 1989); values from Marine ultramicrobacterium strain RB2256 are 0.6 – 3.6E-6 molC/L (Schut *et al.*, 1995). 1.5E-9 and 1.5E-6 molC/L are the calculated values from experimental data for HTCC7211 and HOT1A3, respectively, from non-linear regression, fitting the Michaelis-Menten equation to uptake data.
[i] ~$Q_{INTR}$ when $V_{tran,LALA}$ is half of $V_{max}$ for HTCC7211; range of concentrations used to calibrate were half of the concentration at 0.5 min and concentration at 1 min. Calibration parameter.
[j] Calibration parameter. Set to 10 for HOT1A3 so the term equals 1, for no regulation.
[k] From literature (Grimshaw and Cleland, 1981); value from *B. subtilis* alanine dehydrogenase, 58 molC/gALDM/d; rate was scaled to usual growth rates or $k_{synt}$ of HTCC7211 and HOT1A3 using a growth rate of 8.32 1/d for *B. subtilis*, resulting in a scaling factor of 0.05 – 0.11 and 1.3 – 3.5 for HTCC7211 and HOT1A3, respectively.
[l] From literature (Ohshima *et al.*, 1990; Smith and Emerich, 1993); values ranged from 1.48E-4 – 4.2E-3 molC/L cytoplasm; values were calculated assuming cell volume is 70% cytoplasm in HOT1A3 based on E. coli, and 63.3% in HTCC7211 (Cayley *et al.*, 1991; Zhao *et al.*, 2017)

[m] From literature (Lüderitz and Klemme, 1977; Kresze and Ronft, 1981); values ranged from 1.5 – 23.5 IU/mg protein; growth rates of 4.2 and 9.6 1/d were used for *Rhodospirillum rubrum* and baker's yeast, respectively, to scale the values by growth rate to HYTCC7211 and HOT1A3.

[n] From literature (Visser *et al.*, 1980; Snoep *et al.*, 1992); values ranged from 8.2E-5 to 4.5E-4 molC/L cytoplasm

[o] From literature (Dennis and Bremer, 2008); values ranged from 4.3E5 to 23.4E5 nt/min/cell depending on growth rate. Values were scaled by growth rate of species for the model.

[p] ~$Q_{INTR}$ when $V_{tran,LALA}$ is half of $V_{max}$ for HOT1A3; range of concentrations used to calibrate were from 1 – 5 min time points.

[q] Calibration parameter. Set to 10 so the term equals 1 for HTCC7211, for no regulation.

[r] From literature (Jasin *et al.*, 1985; Francklyn and Schimmel, 1989); values ranged from 8E-8 to 8E-4 molC/L cytoplasm

[s] Calibration parameter

[t] From literature (Hambraeus *et al.*, 2003; Moran *et al.*, 2013); mRNA half-life values ranged from 2 – 7 min. Decay rate calculated using the equation: ln(2)/half-life.

[u] From literature (Koch and Levy, 1955; Moran *et al.*, 2013); protein half-lives ranged from 20 – 70 h. Decay rate calculated using the equation: ln(2)/half-life.

[v] Assumes decay rate of mature, folded ald enzyme is the same as ald protein being translated.

[w] From literature (Kepes, 1963; Kaempfer and Magasanik, 1967); values for maturation time ranged from 1 – 2 min; value calculated is 1/maturation time

[x] ~$Q_{ALDM}$ when fully expressed

[y] Calibration parameter

[z] Calibration parameter; basal uptake is represented as a fraction of the total uptake where uptake is regulated (i.e., in HTCC7211, not in HOT1A3)

[aa] To reflect the starved nature of the cells, initial pyruvate and alanine levels in the cells were set very low.

[ab] From literature (Maier *et al.*, 2011); average value was 0.04 mRNA copies/cell; in HOT1A3, uninduced, we assume ALDT levels are negligible.

[ac] Calibration parameter; at least 10X lower than ALDM

[ad] Calibration parameter; level of mature ald in cells before the start of the experiment; in same range as $f_{ALD/DW}$

**Table S3** Output of mechanistic model for HTCC7211 L-alanine uptake/metabolism under a scenario when cells encounter a patch of L-alanine for 2 minutes, then drift out, with the simulation running until 40 minutes to allow for complete metabolism of any L-alanine encountered. We compared the outcomes for the cells either when they have or lack transcriptional regulation of alanine dehydrogenase. For definitions of the parameters listed and associated units, see Table S2. Calculation of moles ATP produced is based on the assumption of 15 ATP produced per pyruvate molecule.

| | **Maximum value without transcriptional regulation** | **Maximum value with transcriptional regulation** | **$Log_2$ fold change (with / without regulation)** |
|---|---|---|---|
| $Q_{LALA}$ | 2.37E-08 | 1.09E-06 | 5.5 |
| $Q_{PYRU}$ | 5.89E-08 | 7.83E-10 | -6.2 |
| $S_{DIOX}$ | 2.04E-10 | 8.72E-12 | -4.5 |
| $Q_{ALDT}$ | 2.45E-04 | 1.74E-03 | 2.8 |
| $Q_{ALDN}$ | 1.65E-02 | 1.70E-06 | -13.2 |
| $Q_{ALDM}$ | 3.86E-02 | 1.17E-05 | -11.7 |
| $Q_{PROT}$ | 3.08E-04 | 3.02E-03 | 3.3 |
| ATP (moles / gDW) | 2.16E-6 | 2.87E-8 | -6.2 |

**Table S4** List of assemblies used for the bioinformatic comparison of marine oligotrophs and copiotrophs.

Listed predicted maximal growth rates were generated in Weissman et al., 2021.

| Accession | Predicted max growth rate (1/d) | Predicted lifestyle | Genome Name |
|---|---|---|---|
| GCA_002700095.1 | 3.638415297 | Copiotroph | Fulvimarina sp. strain NAT173 |
| GCA_002723235.1 | 5.377013265 | Oligotroph | Spongiibacteraceae bacterium strain SP181 |
| GCA_002711675.1 | 8.083851159 | Oligotroph | Rhodobiaceae bacterium strain SAT202 |
| GCA_002707865.1 | 0.376655473 | Copiotroph | Magnetococcales bacterium strain SAT227 |
| GCA_002715175.1 | 2.728241651 | Copiotroph | Hirschia sp. strain SAT115 |
| GCA_002721695.1 | 12.82181251 | Oligotroph | Roseibacillus sp. strain SP242 |
| GCA_002701545.1 | 14.4781772 | Oligotroph | Blastopirellula sp. strain NAT61 |
| GCA_002694685.1 | 7.420427467 | Oligotroph | Balneola sp. strain IN10 |
| GCA_002693265.1 | 4.532787487 | Copiotroph | Dinoroseobacter sp. strain EAC670 |
| GCA_002700145.1 | 2.572320501 | Copiotroph | Gordonia sp. strain NAT169 |
| GCA_002711645.1 | 3.993858547 | Copiotroph | Desulfovibrio sp. strain SP109 |
| GCA_002723255.1 | 2.687332076 | Copiotroph | Pelagibacterium sp. strain SP179 |
| GCA_002694985.1 | 6.505693706 | Oligotroph | Parvibaculum sp. strain EAC77 |
| GCA_002684995.1 | 1.371741849 | Copiotroph | Oceanospirillum sp. strain CPC24 |
| GCA_002690685.1 | 7.941330113 | Oligotroph | Nisaea sp. strain MED717 |
| GCA_002706335.1 | 3.616477257 | Copiotroph | Ahrensia sp. strain SAT55 |
| GCA_002729995.1 | 1.122881587 | Copiotroph | Rhodobacteraceae bacterium strain NP31 |
| GCA_002724845.1 | 2.910871323 | Copiotroph | Synechococcus sp. RS344 |
| GCA_002687855.1 | 1.921292991 | Copiotroph | Halobacteriovoraceae bacterium strain ARS14 |
| GCA_002700185.1 | 5.215624657 | Oligotroph | Pseudozobellia sp. strain NAT152 |
| GCA_002683255.1 | 2.537477146 | Copiotroph | Oceanicaulis sp. strain CPC78 |
| GCA_002696485.1 | 5.252951463 | Oligotroph | Altibacter sp. strain EAC109 |
| GCA_002688935.1 | 0.568889977 | Copiotroph | Aestuariibacter sp. strain ARS1024 |
| GCA_002717045.1 | 11.54289887 | Oligotroph | Opitutaceae bacterium strain SP4023 |
| GCA_002712385.1 | 3.615556871 | Copiotroph | Bdellovibrionaceae bacterium strain SAT16 |
| GCA_002683755.1 | 1.999571485 | Copiotroph | Sphingorhabdus sp. strain CPC54 |
| GCA_002713165.1 | 9.906837579 | Oligotroph | Verrucomicrobia bacterium strain SAT140 |
| GCA_002726235.1 | 13.42719039 | Oligotroph | Nitrospiraceae bacterium strain NP85 |
| GCA_002712625.1 | 7.200596218 | Oligotroph | Saprospirales bacterium strain SAT153 |
| GCA_002697245.1 | 1.388509188 | Copiotroph | Thioclava sp. strain NAT82 |
| GCA_002699205.1 | 6.927808309 | Oligotroph | Muricauda sp. strain NAT111 |
| GCA_002700365.1 | 5.588324531 | Oligotroph | Kordiimonas sp. strain NAT158 |
| GCA_002712525.1 | 10.65170006 | Oligotroph | Pelagibacteraceae bacterium strain SAT1561 |
| GCA_002716325.1 | 11.11754575 | Oligotroph | Solibacterales bacterium strain SP4380 |
| GCA_002729225.1 | 8.748107241 | Oligotroph | Nitrospinae bacterium strain NP50 |
| GCA_002697225.1 | 11.78375577 | Oligotroph | Candidatus Poribacteria bacterium strain NAT81 |
| GCA_002685435.1 | 2.945650252 | Copiotroph | Cryomorphaceae bacterium strain CPC1326 |
| GCA_002684175.1 | 13.80458105 | Oligotroph | Synechococcus sp. CPC35 |
| GCA_002706715.1 | 2.519817527 | Copiotroph | Synechococcus sp. SAT82 |

| | | | |
|---|---|---|---|
| GCA_002699105.1 | 5.04591367 | Oligotroph | Ectothiorhodospiraceae bacterium strain NAT122 |
| GCA_002699795.1 | 9.485266373 | Oligotroph | Pelagibacterales bacterium strain MED965 |
| GCA_002689785.1 | 10.57139902 | Oligotroph | Prochlorococcus sp. MED630 |
| GCA_002714225.1 | 5.499899528 | Oligotroph | Acidimicrobiaceae bacterium strain SAT1333 |
| GCA_002708205.1 | 12.89467244 | Oligotroph | Gammaproteobacteria bacterium strain SAT2910 |
| GCA_002722295.1 | 3.414728034 | Copiotroph | Stappia sp. strain SP245 |
| GCA_002698505.1 | 4.722592751 | Copiotroph | Synechococcus sp. NAT40 |
| GCA_002684355.1 | 10.76265525 | Oligotroph | Formosa sp. strain CPC323 |
| GCA_002719495.1 | 1.420570007 | Copiotroph | Altererythrobacter sp. strain SP3021 |
| GCA_002692445.1 | 6.221233596 | Oligotroph | Proteobacteria bacterium strain MED1036 |
| GCA_002738205.1 | 5.356655352 | Oligotroph | Hahellaceae bacterium strain EAC91 |
| GCA_002708395.1 | 7.290734609 | Oligotroph | Dehalococcoidia bacterium strain SAT2604 |
| GCA_002713685.1 | 10.48307925 | Oligotroph | Opitutae bacterium strain SAT156 |
| GCA_002712175.1 | 3.829596922 | Copiotroph | Roseovarius sp. strain SAT174 |
| GCA_002683615.1 | 2.770078598 | Copiotroph | Hyphomonas sp. strain CPC7 |
| GCA_002687435.1 | 16.59965732 | Oligotroph | Desulfobacter sp. strain ARS36 |
| GCA_002702105.1 | 8.111550394 | Oligotroph | Nitrosomonadales bacterium strain NAT283 |
| GCA_002707605.1 | 2.565665176 | Copiotroph | Hoeflea sp. strain SAT27 |
| GCA_002729495.1 | 11.38183005 | Oligotroph | Chromatiales bacterium strain NP37 |
| GCA_002683575.1 | 1.31710519 | Copiotroph | Bermanella sp. strain CPC64 |
| GCA_002694165.1 | 4.216702248 | Copiotroph | Alcaligenaceae bacterium strain IN38 |
| GCA_002726575.1 | 14.0447342 | Oligotroph | Porticoccaceae bacterium strain NP64 |
| GCA_002709775.1 | 0.473051457 | Copiotroph | Thalassospira sp. strain SP131 |
| GCA_002731895.1 | 7.483892499 | Oligotroph | Cytophagia bacterium strain NP1152 |
| GCA_002693985.1 | 3.008668181 | Copiotroph | Phenylobacterium sp. strain IN8 |
| GCA_002704865.1 | 1.089778798 | Copiotroph | Sphingobium sp. strain NAT216 |
| GCA_002689065.1 | 8.912136695 | Oligotroph | Puniceicoccaceae bacterium strain ARS1008 |
| GCA_002724995.1 | 11.39181686 | Oligotroph | Anaerolineaceae bacterium strain NP995 |
| GCA_002710305.1 | 4.863991787 | Copiotroph | Nioella sp. strain SAT90 |
| GCA_002687035.1 | 2.448223308 | Copiotroph | Croceicoccus sp. strain ARS62 |
| GCA_002696985.1 | 3.87206859 | Copiotroph | Mesonia sp. strain NAT92 |
| GCA_002714705.1 | 5.072445071 | Oligotroph | Pusillimonas sp. strain SAT110 |
| GCA_002690455.1 | 4.494771484 | Copiotroph | Kiloniella sp. strain MED755 |
| GCA_002729595.1 | 1459.115933 | Oligotroph | Thalassobius sp. strain NP30 |
| GCA_002710155.1 | 2.55249299 | Copiotroph | Halomonadaceae bacterium strain SP1 |
| GCA_002688835.1 | 1.09362471 | Copiotroph | Colwelliaceae bacterium strain ARS1043 |
| GCA_002706085.1 | 1.774915093 | Copiotroph | Alcanivoracaceae bacterium strain SAT93 |
| GCA_002731975.1 | 14.15821072 | Oligotroph | Coxiellaceae bacterium strain NP1046 |
| GCA_002694085.1 | 18.11709065 | Oligotroph | Marinovum sp. strain IN44 |
| GCA_002719735.1 | 3.005292463 | Copiotroph | Alcanivorax sp. strain sp32 |
| GCA_002708995.1 | 3.433586256 | Copiotroph | Geminicoccus sp. strain SP156 |
| GCA_002691585.1 | 8.152056621 | Oligotroph | Legionellales bacterium strain MED607 |
| GCA_002705105.1 | 18.74514448 | Oligotroph | Alteromonas sp. strain NAT174 |
| GCA_002721975.1 | 8.71177666 | Oligotroph | Epsilonproteobacteria bacterium strain SP22 |
| GCA_002694845.1 | 1.896877392 | Copiotroph | Tistrella sp. strain EAC87 |

| | | | |
|---|---|---|---|
| GCA_002717505.1 | 9.119550638 | Oligotroph | Leptospiraceae bacterium strain SP332 |
| GCA_002726835.1 | 6.120637736 | Oligotroph | Haliea sp. strain NP9 |
| GCA_002691945.1 | 0.643349317 | Copiotroph | Cyanobium sp. MED195 |
| GCA_002698345.1 | 11.1443406 | Oligotroph | Chloroflexi bacterium strain NAT441 |
| GCA_002706965.1 | 1.267002582 | Copiotroph | Pseudoalteromonas sp. strain SAT69 |
| GCA_002729155.1 | 3.436198358 | Copiotroph | Leifsonia sp. strain NP56 |
| GCA_002714575.1 | 3.61322229 | Copiotroph | Flavobacterium sp. strain SAT130 |
| GCA_002686705.1 | 13.41329344 | Oligotroph | Bacteroidetes bacterium strain ARS83 |
| GCA_002699145.1 | 9.461316289 | Oligotroph | Halioglobus sp. strain NAT121 |
| GCA_002729885.1 | 9.834001409 | Oligotroph | Spirochaetaceae bacterium strain NP47 |
| GCA_002688185.1 | 11.78061134 | Oligotroph | Spirochaetales bacterium strain ARS1246 |
| GCA_002684335.1 | 9.738473262 | Oligotroph | Candidatus Puniceispirillum sp. strain CPC325 |
| GCA_002708475.1 | 2.756113149 | Copiotroph | Oleispira sp. strain SAT217 |
| GCA_002683145.1 | 4.009117584 | Copiotroph | Myxococcales bacterium strain CPC88 |
| GCA_002724745.1 | 0.831545355 | Copiotroph | Herbaspirillum sp. strain RS355 |
| GCA_002706985.1 | 8.891782081 | Oligotroph | Sneathiella sp. strain SAT66 |
| GCA_002699365.1 | 6.192410144 | Oligotroph | Coraliomargarita sp. strain NAT145 |
| GCA_002687285.1 | 1.19733733 | Copiotroph | Pelagibaca sp. strain ARS5 |
| GCA_002721535.1 | 6.06000897 | Oligotroph | Oceanospirillaceae bacterium strain SP264 |
| GCA_002701745.1 | 4.075326804 | Copiotroph | Flammeovirgaceae bacterium strain NAT44 |
| GCA_002727935.1 | 11.36259664 | Oligotroph | Dehalococcoidaceae bacterium strain RS451 |
| GCA_002700765.1 | 4.320978634 | Copiotroph | Synechococcus sp. MED850 |
| GCA_002689605.1 | 8.056774889 | Oligotroph | Filomicrobium sp. strain MED665 |
| GCA_002715485.1 | 5.02833414 | Oligotroph | Aequorivita sp. strain SP75 |
| GCA_002710765.1 | 5.877304406 | Oligotroph | Rickettsiales bacterium strain SAT45 |
| GCA_002683405.1 | 23.73489494 | Oligotroph | Salinisphaeraceae bacterium strain CPC72 |
| GCA_002708825.1 | 3.903963892 | Copiotroph | Phyllobacteriaceae bacterium strain SP17 |
| GCA_002687555.1 | 2.327891558 | Copiotroph | Blastomonas sp. strain ARS3 |
| GCA_002702685.1 | 6.024286661 | Oligotroph | Planctomyces sp. strain NAT128 |
| GCA_002695065.1 | 2.916753762 | Copiotroph | Salinicola sp. strain EAC73 |
| GCA_002701665.2 | 3.543256062 | Copiotroph | Actinomycetales bacterium strain NAT49 |
| GCA_002706905.1 | 5.178884314 | Oligotroph | Pseudooceanicola sp. strain SAT54 |
| GCA_002684555.1 | 8.532380562 | Oligotroph | Gimesia sp. strain CPC31 |
| GCA_002693185.1 | 1.603979161 | Copiotroph | Gramella sp. strain EAC68 |
| GCA_002716765.1 | 9.279894626 | Oligotroph | Candidatus Magasanikbacteria bacterium strain SP4150 |
| GCA_002693945.1 | 1.657717047 | Copiotroph | Oleibacter sp. strain IN912 |
| GCA_002686465.1 | 16.33467873 | Oligotroph | Candidatus Woesebacteria bacterium strain ARS1183 |
| GCA_002703405.1 | 1.75507211 | Copiotroph | Oceanibulbus sp. strain EAC16 |
| GCA_002690325.1 | 11.08910497 | Oligotroph | Synechococcus sp. ARS1019 |
| GCA_002699025.1 | 3.860081671 | Copiotroph | Sandaracinus sp. strain NAT131 |
| GCA_002694065.1 | 21.83200753 | Oligotroph | Acidiferrobacter sp. strain IN47 |
| GCA_002725135.1 | 9.613298221 | Oligotroph | Actinobacteria bacterium strain NP978 |
| GCA_002693695.1 | 6.311313467 | Oligotroph | Pseudohongiella sp. strain EAC47 |
| GCA_002688135.1 | 11.5419686 | Oligotroph | Parcubacteria group bacterium strain ARS125 |
| GCA_002711735.1 | 6.160415282 | Oligotroph | Ilumatobacter sp. strain SAT196 |

| | | | |
|---|---|---|---|
| GCA_002693505.1 | 4.229975364 | Copiotroph | Hyphomonadaceae bacterium strain EAC632 |
| GCA_002694425.1 | 3.615344842 | Copiotroph | Arenimonas sp. strain IN24 |
| GCA_002705125.1 | 10.96325055 | Oligotroph | Verrucomicrobiales bacterium strain NAT181 |
| GCA_002725975.1 | 1.13289652 | Copiotroph | Pseudomonadaceae bacterium strain NP95 |
| GCA_002698575.1 | 2.528104209 | Copiotroph | Pimelobacter sp. strain NAT253 |
| GCA_002706995.1 | 2.308943865 | Copiotroph | Aurantimonas sp. strain SAT5 |
| GCA_002713615.1 | 6.552634153 | Oligotroph | SAR116 cluster bacterium strain SAT158 |
| GCA_002721925.1 | 10.62601893 | Oligotroph | Micrococcales bacterium strain SP226 |
| GCA_002726755.1 | 10.94348441 | Oligotroph | Flavobacteriales bacterium strain NP93 |
| GCA_002722955.1 | 14.99194545 | Oligotroph | Planctomycetaceae bacterium strain SP200 |
| GCA_002724575.1 | 14.83538558 | Oligotroph | Gemmatimonadetes bacterium strain RS373 |
| GCA_002726815.1 | 4.337535038 | Copiotroph | Methylophaga sp. strain NP92 |
| GCA_002706265.1 | 5.943540639 | Oligotroph | Polycyclovorans sp. strain SAT60 |
| GCA_002686115.1 | 7.015029234 | Oligotroph | Halobacteriovorax sp. strain ARS37 |
| GCA_002732075.1 | 5.186404421 | Oligotroph | Nevskiales bacterium strain NP100 |
| GCA_002710685.1 | 3.703860564 | Copiotroph | Cytophagaceae bacterium strain SAT51 |
| GCA_002700895.1 | 2.776827944 | Copiotroph | Cyanobium sp. MED843 |
| GCA_002711755.1 | 3.270437239 | Copiotroph | Confluentimicrobium sp. strain SAT198 |
| GCA_002707795.1 | 2.936488346 | Copiotroph | Maricaulis sp. strain SAT23 |
| GCA_002715525.1 | 5.629290331 | Oligotroph | Owenweeksia sp. strain SP74 |
| GCA_002697475.1 | 4.699000549 | Copiotroph | Aquimarina sp. strain NAT579 |
| GCA_002695375.1 | 2.687087058 | Copiotroph | Rhizobiaceae bacterium strain EAC683 |
| GCA_002690755.1 | 4.71753351 | Copiotroph | Phycisphaerae bacterium strain MED708 |
| GCA_002684455.1 | 3.33787998 | Copiotroph | Arcobacter sp. strain CPC309 |
| GCA_002707785.1 | 2.11287515 | Copiotroph | Cellvibrionaceae bacterium strain SAT230 |
| GCA_002728955.1 | 3.079652448 | Copiotroph | Cyanobium sp. RS427 |
| GCA_002687115.1 | 9.630169983 | Oligotroph | Cyanobium sp. ARS6 |
| GCA_002687495.1 | 1.287954175 | Copiotroph | Micavibrio sp. strain ARS33 |
| GCA_002693125.1 | 0.77581447 | Copiotroph | Roseobacter sp. strain EAC695 |
| GCA_002720205.1 | 8.646581597 | Oligotroph | Prochlorococcus sp. SP3034 |
| GCA_002704615.1 | 13.68947321 | Oligotroph | Cellvibrionales bacterium strain MED776 |
| GCA_002714685.1 | 0.736983154 | Copiotroph | Cobetia sp. strain SAT113 |
| GCA_002722255.1 | 3.890162666 | Copiotroph | Winogradskyella sp. strain SP247 |
| GCA_002694415.1 | 2.673400752 | Copiotroph | Citromicrobium sp. strain IN20 |
| GCA_002704915.1 | 6.857291378 | Oligotroph | Parvularcula sp. strain NAT21 |
| GCA_002686945.1 | 8.778094845 | Oligotroph | Planctomycetes bacterium strain ARS68 |
| GCA_002716305.1 | 1.219413721 | Copiotroph | Peredibacter sp. strain SP55 |
| GCA_002705365.1 | 14.98960404 | Oligotroph | Candidatus Marinimicrobia bacterium strain MED806 |
| GCA_002708795.1 | 6.13370507 | Oligotroph | Paracoccus sp. strain SP169 |
| GCA_002698995.1 | 5.901587144 | Oligotroph | Ignavibacteriae bacterium strain NAT134 |
| GCA_002719855.1 | 7.555211648 | Oligotroph | Legionellaceae bacterium strain SP3112 |
| GCA_002685455.1 | 10.37466362 | Oligotroph | Porticoccus sp. strain CPC1232 |
| GCA_002730815.1 | 1.798274385 | Copiotroph | Spongiibacter sp. strain NP125 |
| GCA_002721515.1 | 15.19353421 | Oligotroph | Nitrospina sp. strain SP265 |
| GCA_002727395.1 | 11.54279429 | Oligotroph | Rhodospirillales bacterium strain RS665 |

| | | | |
|---|---|---|---|
| GCA_002729215.1 | 10.25789774 | Oligotroph | Rubrivirga sp. strain NP52 |
| GCA_002685495.1 | 10.95129627 | Oligotroph | Woeseiaceae bacterium strain CPC1328 |
| GCA_002702445.1 | 7.254501592 | Oligotroph | Propionibacteriaceae bacterium strain NAT249 |
| GCA_002707295.1 | 8.02637144 | Oligotroph | Flavobacteriaceae bacterium strain SAT2981 |
| GCA_002725655.1 | 14.08208274 | Oligotroph | Opitutales bacterium strain NP990 |
| GCA_002721595.1 | 2.437456046 | Copiotroph | Idiomarinaceae bacterium strain SP259 |
| GCA_002692105.1 | 10.5380197 | Oligotroph | Marinoscillum sp. strain MED1001 |
| GCA_002720595.1 | 16.47191358 | Oligotroph | Woeseia sp. strain SP2978 |
| GCA_002720855.1 | 11.56102693 | Oligotroph | Rhodospirillaceae bacterium strain SP3 |
| GCA_002697745.1 | 3.909644412 | Copiotroph | Nitratireductor sp. strain NAT291 |
| GCA_002684595.1 | 1.386951466 | Copiotroph | Novosphingobium sp. strain CPC302 |
| GCA_002710665.1 | 3.169201128 | Copiotroph | Sutterellaceae bacterium strain SAT62 |
| GCA_002708315.1 | 8.038152594 | Oligotroph | Lentimicrobiaceae bacterium strain SAT2766 |
| GCA_002695005.1 | 3.467737238 | Copiotroph | Maritimibacter sp. strain EAC76 |
| GCA_002725875.1 | 19.86785733 | Oligotroph | Deltaproteobacteria bacterium strain NP956 |
| GCA_002686735.1 | 1.85098234 | Copiotroph | Psychrobacter sp. strain ARS82 |
| GCA_002690525.1 | 6.766651249 | Oligotroph | SAR324 cluster bacterium strain MED745 |
| GCA_002726335.1 | 13.44948736 | Oligotroph | Gemmatimonadaceae bacterium strain NP81 |
| GCA_002704825.1 | 1.469955085 | Copiotroph | Phycisphaeraceae bacterium strain NAT22 |
| GCA_002720235.1 | 3.654066073 | Copiotroph | Henriciella sp. strain SP3031 |
| GCA_002707985.1 | 2.52158417 | Copiotroph | Sphingomonas sp. strain SAT22 |
| GCA_002706605.1 | 11.75794406 | Oligotroph | Rhodovulum sp. strain SAT37 |
| GCA_002727475.1 | 9.121443385 | Oligotroph | Thiotrichales bacterium strain RS615 |
| GCA_002683935.1 | 1.139587973 | Copiotroph | Erythrobacter sp. strain CPC47 |
| GCA_002697445.1 | 4.774742336 | Copiotroph | Rhodothermaceae bacterium strain NAT6 |
| GCA_002718235.1 | 4.451400674 | Copiotroph | Oceanospirillales bacterium strain SP333 |
| GCA_002693285.1 | 2.599940083 | Copiotroph | Synechococcus sp. EAC657 |
| GCA_002691215.1 | 2.66204894 | Copiotroph | Ponticaulis sp. strain MED653 |
| GCA_002691025.1 | 14.45378951 | Oligotroph | Spirochaeta sp. strain MED672 |
| GCA_002707725.1 | 4.821988212 | Copiotroph | Kangiellaceae bacterium strain SAT2587 |
| GCA_002729445.1 | 1.419584562 | Copiotroph | Variovorax sp. strain NP4 |
| GCA_002715985.1 | 1363.950729 | Oligotroph | Xanthomonadales bacterium strain SP48 |
| GCA_002699305.1 | 2.145630606 | Copiotroph | Martelella sp. strain NAT10 |
| GCA_002714405.1 | 3.190875994 | Copiotroph | Cyanobium sp. SAT1300 |
| GCA_002683675.1 | 0.834876353 | Copiotroph | Rheinheimera sp. strain CPC6 |
| GCA_002706925.1 | 3.160067018 | Copiotroph | Rhizobiales bacterium strain SAT53 |
| GCA_002707365.1 | 17.22543865 | Oligotroph | Trueperaceae bacterium strain SAT2911 |
| GCA_002729635.1 | 3.710228221 | Copiotroph | Mameliella sp. strain NP29 |
| GCA_002703585.1 | 3.855598523 | Copiotroph | Halieaceae bacterium strain EAC26 |
| GCA_002727035.1 | 2.229825313 | Copiotroph | Methylobacterium sp. strain NP6 |
| GCA_002699035.1 | 10.19678033 | Oligotroph | Verrucomicrobiaceae bacterium strain NAT132 |
| GCA_002730245.1 | 9.828809328 | Oligotroph | Candidatus Pelagibacter sp. strain NP144 |
| GCA_002700385.1 | 7.227395319 | Oligotroph | Cycloclasticus sp. strain NAT157 |
| GCA_002692385.1 | 6.585314634 | Oligotroph | Prochlorococcus sp. MED105 |
| GCA_002698305.1 | 4.344154348 | Copiotroph | Candidatus Campbellbacteria bacterium strain NAT484 |

| | | | |
|---|---|---|---|
| GCA_002691345.1 | 4.621866613 | Copiotroph | Synechococcus sp. MED650 |
| GCA_002731235.1 | 1.168258383 | Copiotroph | Pseudoalteromonadaceae bacterium strain NP121 |
| GCA_002701375.1 | 4.603030047 | Copiotroph | Cyanobium sp. NAT70 |
| GCA_002690705.1 | 14.83131151 | Oligotroph | Magnetovibrio sp. strain MED |
| GCA_002729835.1 | 6.152335804 | Oligotroph | Synechococcus sp. NP17 |
| GCA_002713265.1 | 16.04858896 | Oligotroph | Acidobacteria bacterium strain SAT1391 |
| GCA_002729955.1 | 5.128587217 | Oligotroph | Salinisphaera sp. strain NP40 |
| GCA_002698165.1 | 8.537079162 | Oligotroph | Planktomarina sp. strain NAT682 |
| GCA_002691095.1 | 3.698824127 | Copiotroph | Kiritimatiellaceae bacterium strain MED670 |
| GCA_002685105.1 | 1.66320046 | Copiotroph | Pseudomonadales bacterium strain CPC18 |
| GCA_002721545.1 | 9.017957506 | Oligotroph | Nitrosomonadaceae bacterium strain SP263 |
| GCA_002692525.1 | 1.569598362 | Copiotroph | Pseudomonas sp. strain IN922 |
| GCA_002732055.1 | 1.686734378 | Copiotroph | Halomonas sp. strain NP1 |
| GCA_002716065.1 | 6.912202552 | Oligotroph | Crocinitomicaceae bacterium strain SP82 |
| GCA_002693245.1 | 13.16476267 | Oligotroph | Acidiferrobacteraceae bacterium strain EAC671 |
| GCA_002699225.1 | 2.444849084 | Copiotroph | Candidatus Saccharibacteria bacterium strain NAT105 |
| GCA_002686885.1 | 12.16229433 | Oligotroph | Pedosphaera sp. strain ARS72 |
| GCA_002693615.1 | 1.368210504 | Copiotroph | Nocardioides sp. strain EAC54 |
| GCA_002709385.1 | 5.684323161 | Oligotroph | Waddliaceae bacterium strain SP13 |
| GCA_002695745.2 | 6.424715128 | Oligotroph | Rhodobacterales bacterium strain EAC638 |
| GCA_002689885.1 | 6.713980001 | Oligotroph | Candidatus Thioglobus sp. strain MED612 |
| GCA_002731545.1 | 9.601540879 | Oligotroph | Rubinisphaera sp. strain NP109 |
| GCA_002685515.1 | 3.446504546 | Copiotroph | Synechococcus sp. CPC100 |
| GCA_002701555.1 | 5.922600665 | Oligotroph | Alphaproteobacteria bacterium strain NAT608 |
| GCA_002716005.1 | 2.052062898 | Copiotroph | Leeuwenhoekiella sp. strain SP45 |
| GCA_002696855.1 | 0.969290361 | Copiotroph | Alteromonadaceae bacterium strain EAC17 |
| GCA_002694775.1 | 1.691304852 | Copiotroph | Sphingomonadaceae bacterium strain EAC97 |
| GCA_002730395.1 | 10.60139477 | Oligotroph | Dehalococcoidales bacterium strain NP138 |
| GCA_002695825.1 | 4.674701417 | Copiotroph | Algoriphagus sp. strain EAC63 |
| GCA_002696915.1 | 5.39962477 | Oligotroph | Rhodopirellula sp. strain EAC14 |
| GCA_002697045.1 | 2.229570372 | Copiotroph | Erythrobacteraceae bacterium strain EAC1 |
| GCA_002699245.1 | 8.047477395 | Oligotroph | Aestuariivita sp. strain NAT114 |

**Table S5** Primer sequences used in RT-qPCR assays.

| Strain | Gene name | GenBank accession number | Primer sequence | Forward or reverse | Calculated $T_M$ (C) | Amplicon size (bp) |
|---|---|---|---|---|---|---|
| HOT1A3 | *recA* | AMN10922.1 | AAAAGCTTTAACCGCCGCAG | Forward | 66.1 | 165 |
| | *recA* | | ACAACTCGTCCACATGGCAA | Reverse | 65.4 | |
| | *rpoD* | AMN10687 | ACACCGCTATCGTCTTCGTC | Forward | 63.1 | 154 |
| | *rpoD* | | TGAGCGACATCATCTCTGGC | Reverse | 65.5 | |
| | *ald* | AMN11997.1 | ACGCAAACAAAGCTGCTTGT | Forward | 63.7 | 150 |
| | *ald* | | TTTTTCAAGGTAGTGCGCGC | Reverse | 66.1 | |
| HTCC 7211 | *recA* | EDZ60796.1 | AGCTGGAGGAATTTGTGCGT | Forward | 65.5 | 196 |
| | *recA* | | GCGTTAGTGCTGCAACTGAG | Reverse | 64.6 | |
| | *rpoD* | EDZ60035.1 | TACAGGACCCAGTGCAAAGC | Forward | 65.5 | 98 |
| | *rpoD* | | GCGGCAAGCGTTGGATTAAA | Reverse | 65.0 | |
| | *ald* | EDZ60221.1 | GCTGTTGCTGGTCGAATGTC | Forward | 64.6 | 98 |
| | *ald* | | ATCAACTCCTGGTGCTCCAC | Reverse | 65.0 | |

**Table S6** Model parameters for the estimation of patch residence time model.

| Parameter | Units | Value | Notes |
|---|---|---|---|
| $D_{DOM}$ | $m^2/s$ | 3.2e-11 | = 1e-12 – 1e-9 (a)<br>= 3.2e-12 – 3.2e-9 (b)<br>= 1e-10 (c)<br>= 1e-9 (d)<br>= 1e-9 (e) |
| $D$ (motile cell) | $m^2/s$ | 4.7e-13 (f1) | = 4.6e-13, *Vibrio* Stationary 13B01 (f)<br>= 2.5e-13, *Vibrio* Exponential 13B01 (f) |
| $D$ (non-motile cell) | $m^2/s$ | 1.0e-12 (g1) | = 1.1e-12, *Pelagibacter* HTCC1062 (g)<br>= 9.8e-13, *Pelagibacter* HTCC7211 (g) |
| $K_D$ | µmol/L | 10 | = 100 (d)<br>= 3 – 1,000 (e)<br>= 10, 1-10 (h) |
| $\tau_0$ | s | 0.3 (i1) | = 0.67 (d) |
| $T_m$ | s | 0.1 | = 1 (d)<br>= 0.1, 0.1-1 (h) |
| $\alpha$ | s | 30 | = 660 (d)<br>= 30 (h) |
| $D_r$ | $rad^2/s$ | 0.035 | = 0 (d)<br>= 0.035 (h) |
| $v_{mot,ave}$ | µm/s | 40 | $v_{ave}$:<br>= 12 (d)<br>= 12-80 (c)<br>= 36 (h) |
| $v_{mot,logvar}$ | - | 0.20 | = 0.21 (h) |
| $v_{mot,min}$ | µm/s | 5 | = 5 (h) |
| $v_{mot,max}$ | µm/s | 150 | = 150 (h) |
| $f_v$ | - | 1.3 | = 1.3 (h) |
| $C_v$ | µmol/L | 0.050 | = 0.050 (h) |
| $c1$ | s | -0.3942 | = -0.3942 (h) |
| $c2$ | s/µm | -0.1285 (i2) | = -0.2019 (h) |
| $c3$ | s | 0.8452 | = 0.8452 (h) |
| $c4$ | µm/s | 29.66 (i2) | = 18.88 (h) |
| $c5$ | - | 0.086 (i1) | = 0.055 (h) |
| $c6$ | - | 1.1 (i1) | = 0.72 (h) |
| $c7$ | s/µm | -0.16 (i2) | = -0.25 (h) |
| $c8$ | µm/s | 57 (i2) | = 36 (h) |

(a) Solutes consumed by marine bacteria, Taylor and Stocker (Taylor and Stocker, 2012)
(b) Molecules with intermediate diffusivities, Smriga et al. (Smriga *et al.*, 2016)
(d) From Figure 3 and Table 1 of Jackson (Jackson, 1987, 19)
(e) Bowen et al. (Bowen *et al.*, 1993)
(f1) Based on cell size 4.7e-15 molC/cell, assumed motile cells are 10 times larger than non-motile cells. See footnote (f2).
(f2) Calculation diffusion coefficient from size as in Lambert et al. (Lambert *et al.*, 2019), assuming spherical shape and T = 25°C.
(f) Calculated from 5.0e-15 molC/cell, *Vibrio* Stationary 13B01, and 3.3e-14 molC/cell, *Vibrio* Exponential 13B01, from Cermak et al. (Cermak *et al.*, 2017), converted assuming 0.4 gC/gDW. See footnote (f2).

(g1) Based on average observed cell size, 4.7e-16 molC/cell. See footnotes (g) and (f2).
(g) Calculated from 4.0e-16 molC/cell, *Pelagibacter* HTCC1062, and 5.3e-16 molC/cell, *Pelagibacter* HTCC7211, from Cermak et al. (Cermak *et al.*, 2017), converted assuming 0.4 gC/gDW. See footnote (f2).
(h) Son et al. (Son *et al.*, 2016). For $v_{max}$ etc. based on fit to 3D/2D-corrected data in Figure 4A of reference.
(i1) Adjusted/calibrated to observations of Son et al. (Son *et al.*, 2016), see text and Supplemental Methods.
(i2) Velocity parameters adjusted by 1.6 to account for unobserved third dimension (see Supplemental Methods).

### Supplemental Figures

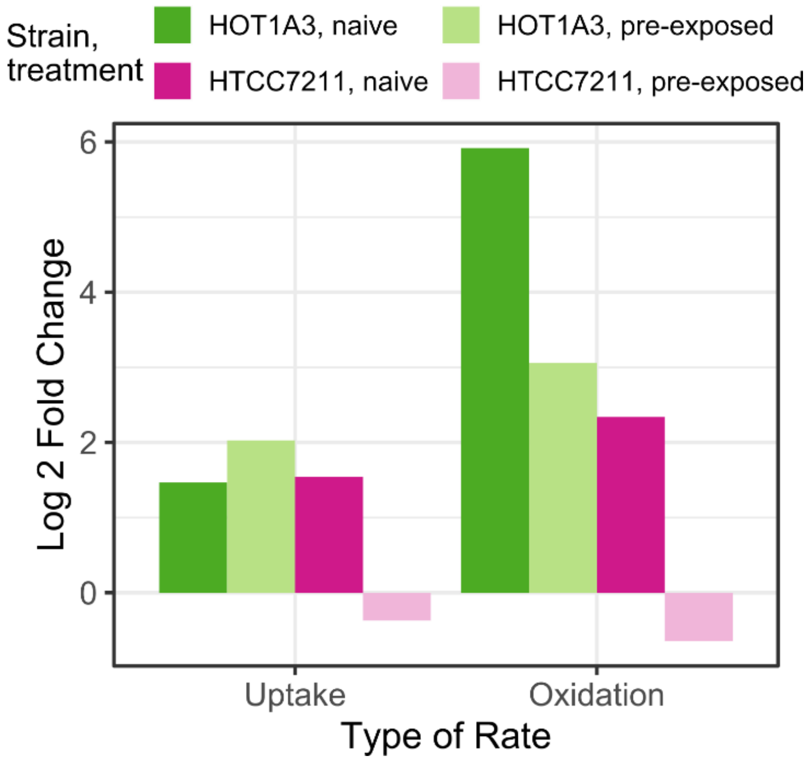


**Figure S1** Comparison of the log 2-fold changes in uptake or oxidation rate of 4 µM [$^{14}$C]L-alanine ([$^{14}$C]LALA) in a model oligotroph, *Ca. P.* st. HTCC7211, and a model copiotroph, *A. macleodii* st. HOT1A3, that were either pre-exposed to 4 µM LALA (lighter colors) or naïve (darker colors). Fold change was calculated as the log 2 change from the initial rate (0 min to the time point just before oxidation initiation; 0 - 0.5 min for HTCC7211 and pre-exposed HOT1A3, 0 – 2 min for naïve HOT1A3) to the final rate (20 – 40 min for naïve cells, 5 - 10 min for pre-exposed cells).

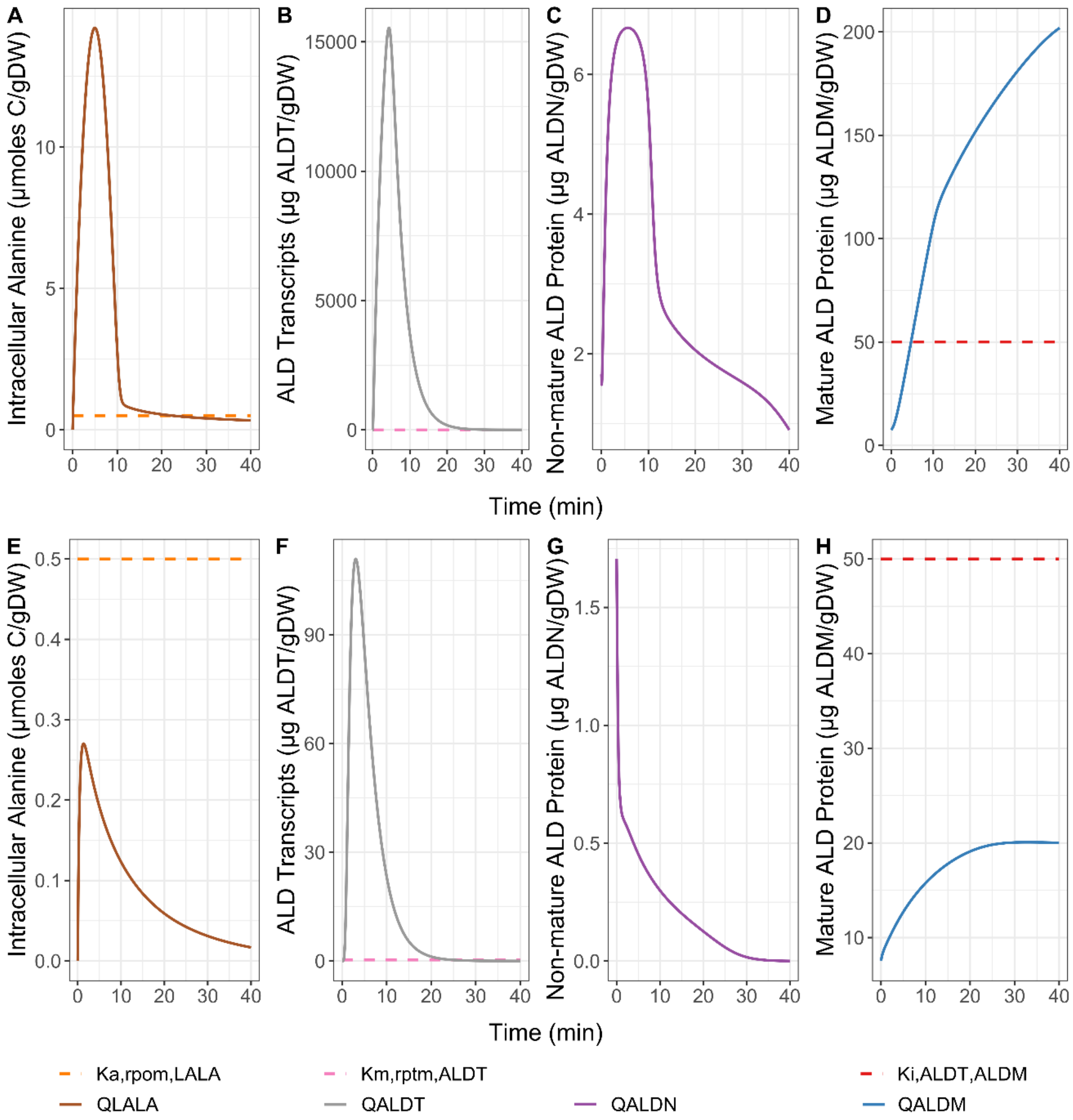


**Figure S2** Output from the mechanistic model to explore various cellular parameters related to the induction of L-alanine (LALA) metabolism by the alanine dehydrogenase enzyme (ALD) in the copiotroph *A. macleodii* st. HOT1A3 at either 4 µM LALA (A-D) or 62.5 nM LALA (E-H). $Q_{LALA}$: intracellular LALA concentration; $K_{a,rpom,LALA}$: activation constant for alanine dehydrogenase transcription; $Q_{ALDT}$: transcript levels for alanine dehydrogenase mRNA; $K_{m,rptm,ALDT}$: half-saturation constant of ribosomes for ALDT; $Q_{ALDN}$: abundance of naïve alanine dehydrogenase enzyme, unfolded; $Q_{ALDM}$: abundance of mature alanine dehydrogenase enzyme; $K_{i,ALDT,ALDM}$: inhibition constant for the inhibition of ALDT production at saturating levels of ALDM. Values for all activation, half-saturation, and inhibition constants are given in Table S2.

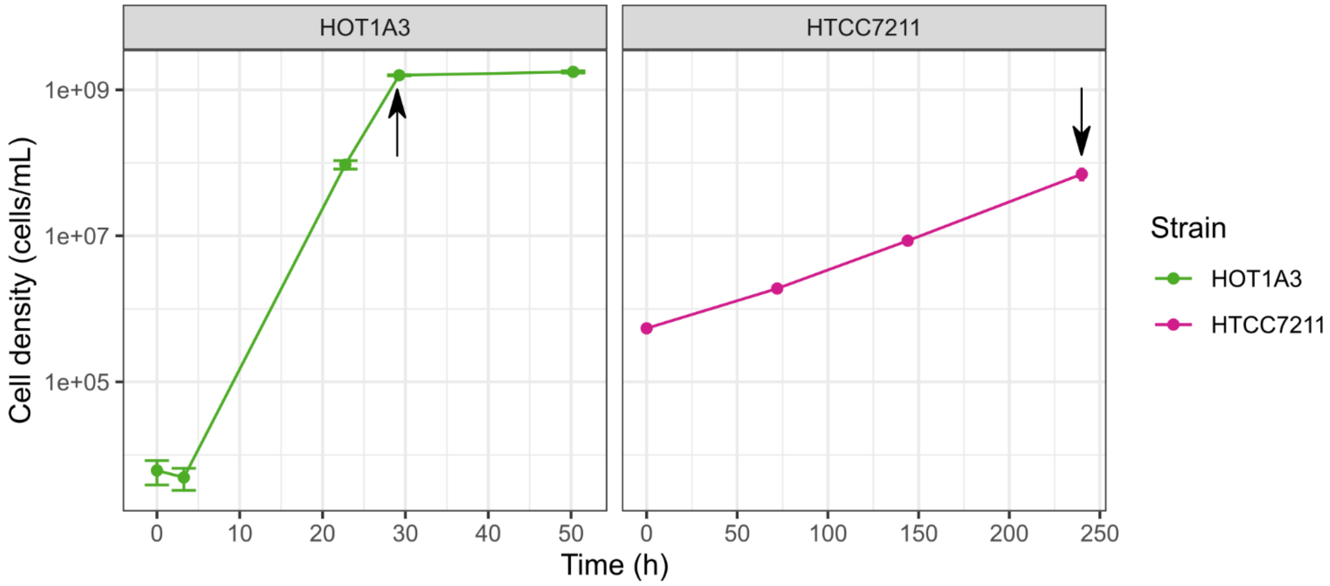


**Figure S3** Representative growth curves for (A) *Alteromonas macleodii* st. HOT1A3 and (B) *Ca.* Pelagibacter st. HTCC7211 cultures used for $^{14}$C-L-alanine uptake experiments. Arrow indicates the time point where cells were harvested for use in experiments. Error bars are standard deviation of duplicate cultures. Where not visible, error bars fall within the size of the data point.

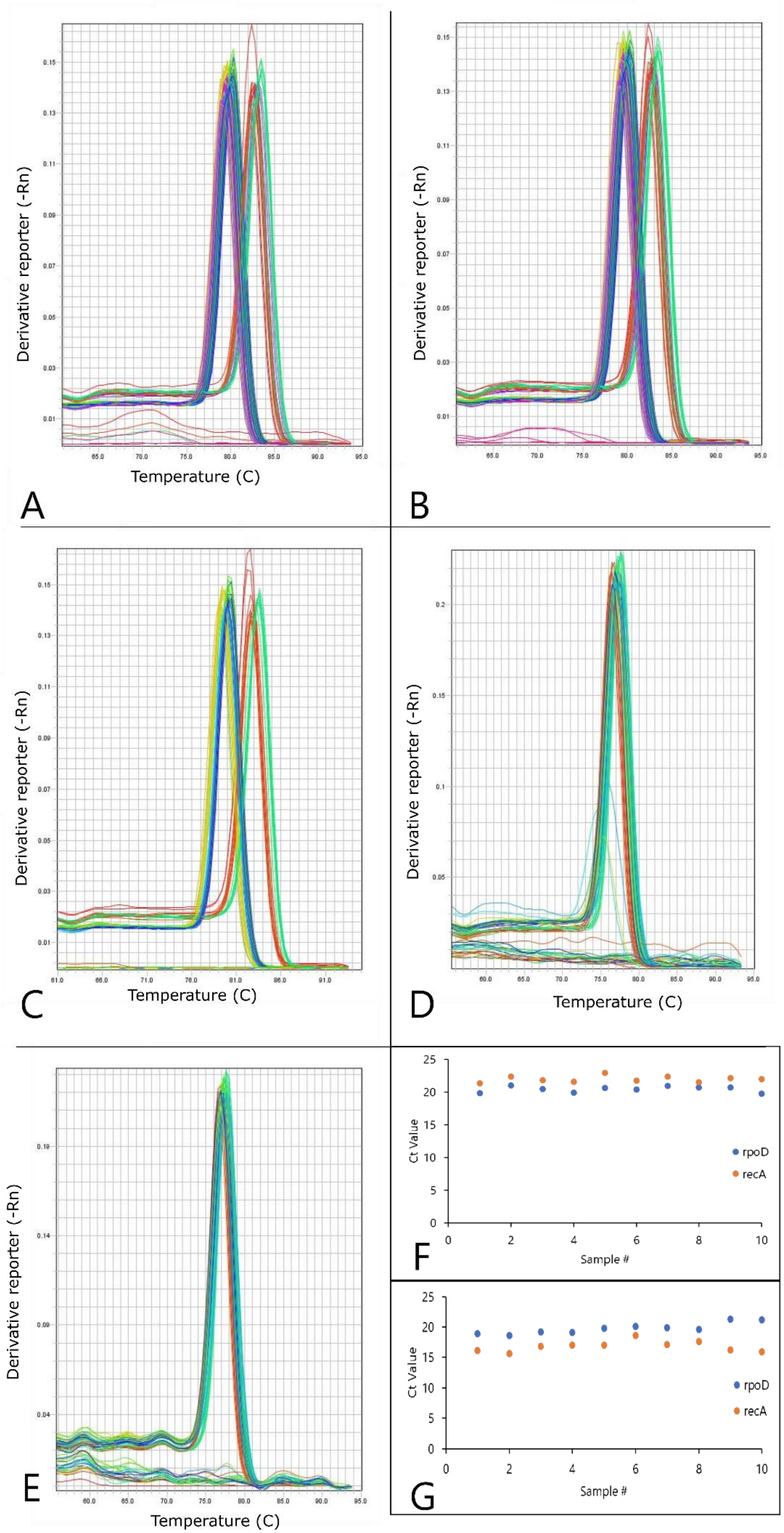

Derivative reporter (-Rn)
Temperature (C)
A
B
C
D
E
Ct Value
Sample #
rpoD
recA
F
G

**Figure S4** Supporting data for RT-qPCR experiments. (A – C) Melt curves from the three HOT1A3 experimental qPCR runs, showing a single peak in all samples for all three genes. Samples amplified by *recA* and *rpoD* had a peak at 80C, while those amplified by *ald* had a peak at 82C. (D – E) Melt curves from the two HTCC7211 experimental qPCR runs, showing a single peak in all samples for all three genes. (F – G) Results of qPCR runs to verify that the endogenous control genes, *recA* and *rpoD*, had consistent expression across all time points and treatments in (F) HOT1A3 and (G) HTCC7211. Values shown are the average of duplicate qPCR runs.

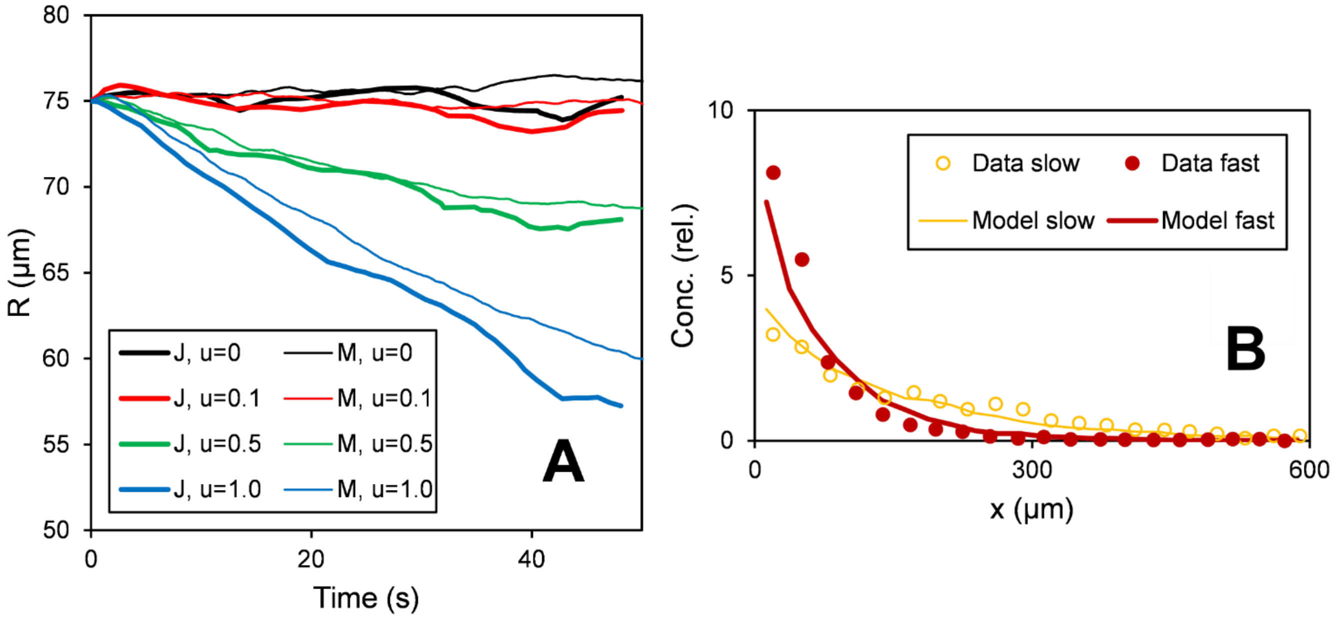


**Figure S5** (A) Internal benchmarking of the run & tumble motility component of the patch simulation model against simulations from Jackson, 1987 (Jackson, 1987). "J" corresponds to Fig. 3 of that paper. "M" corresponds to this model. Small differences between our model and that of Jackson can be attributed to stochasticity. (B) Internal benchmarking of the run-reverse-flick motility component of the patch simulation model against experimental results from (Son *et al.*, 2016).

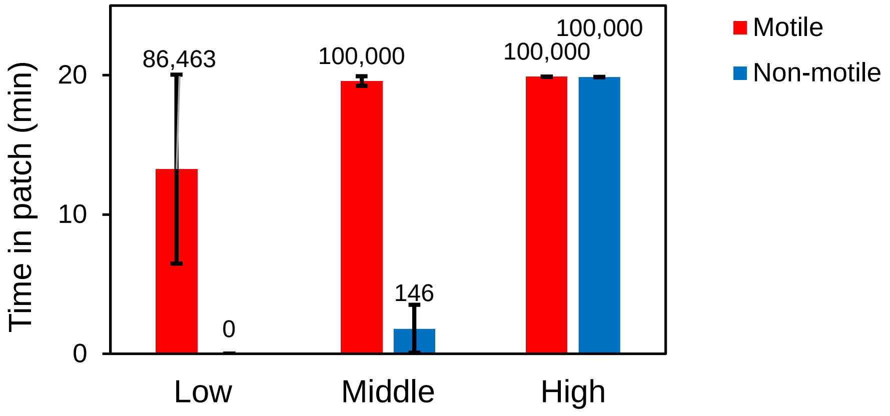


**Figure S6** Exposure time of motile and non-motile cells to a continuous source of nutrients as a function of source size (Middle concentration is the same as in Fig. 5B, Low = Middle / 10, High = Middle × 10). Cells were placed Middle distance from the source, same as in Fig. 5.
